# Conservation of a lateralized visuo-motor axis in hawkmoth proboscis probing

**DOI:** 10.64898/2026.01.27.702022

**Authors:** Lochlan Walsh, Sören Kannegieser, Anna Stöckl

## Abstract

Lateralization of behaviour, including motor control and sensory processing, is widespread across bilaterians. In visually guided tasks it often manifests as an axis aligning eye, appendage, and a target within a shared reference frame, such as eye-hand coordination in humans or eye-beak coordination in birds. While studied intensively in a few vertebrate systems, whether similar control principles apply to invertebrates, and more generally, how sensory and motor lateralization are linked mechanistically, remains unclear.

Using the proboscis inspection behaviour of hummingbird hawkmoth *Macroglossum stellatarum* as a model for visual appendage guidance, our study provides evidence for lateralized visuo-motor control in an invertebrate. Combining high-speed videography and markerless pose estimation, we establish the underlying control axis between the hawkmoths’ unpaired appendage and its eye. We demonstrate that individuals displayed stable, idiosyncratic proboscis lateralization, which was tightly linked to their instantaneous viewing angle of visual targets, thus forming a persistent eye- proboscis-target axis. This axis also had functional consequences for feature-targeting. Assessing the sensory-motor plasticity using monocular occlusion, we found that moths preserved their lateralized visuo-motor geometry by adjusting body posture during flower inspection.

Our findings suggest convergent control principles with vertebrate models of lateralized visual appendage guidance, while highlighting stark differences in sensory-motor plasticity, thus adding to our general understanding of how lateralization shapes control strategies across nervous systems.

**Significance Statement:** Lateralization is widespread across animals and shapes how sensation and action are coordinated. Visually guided reaching with appendages is frequently lateralized across taxa, reflecting biases like handedness and eye dominance. However, the mechanistic link between lateralized sensing and motor control, and the extent of their plasticity, remain poorly understood, particularly in invertebrates. Here we show that the hummingbird hawkmoth integrates individually lateralized vision and proboscis probing movements into a unified control axis. Upon sensory perturbation, moths adjusted their position and body posture to maintain this axis. The visuo-motor lateralization produced measurable functional consequences, revealing it as a form of sensory-motor optimization. These findings uncover convergent principles of lateralized visuo-motor control across insects, humans, birds, and elephants.

## Introduction

Lateralization, expressed in asymmetric motor function, sensation and neural architecture, is a prevalent feature of nervous systems across the animal kingdom (1–4). Lateralization can manifest in sensory pathways as well as at a motor level (5). Visually-guided actions, such as reaching or grasping (6,7), are widely investigated for their sensory-motor asymmetries, where a functional coupling is created between the eye and the appendage. Consistent left-right asymmetries within this coupling constitute visuo-motor lateralization. A typical example is a dominant eye (8) that supports appendage placement, such as eye-hand coordination in humans (9–11) or eye-beak coordination in birds (12–14). Such lateralized guidance forms a visuo-motor axis, where both the target and the moving appendage are kept within the view of one eye (or visual field) (15–18). This simplifies control by aligning perception and action within a shared reference frame (16,19–20). It thus reduces the complexity of sensory-motor transformations involved in tasks such as reaching (6,7,21), potentially even more so when the eye, appendage, and target are on the same side of the body (22). Yet, it remains unclear whether adopting a lateralized visuo-motor axis is a general visual control strategy across taxa. More importantly, whether the sensory or motor components causally drive this asymmetry independently, or reflect an integrated sensory-motor strategy, has rarely been demonstrated (3).

Evidence for lateralized visuo-motor axes comes primarily from vertebrates, ranging from human hand use (10,11,16), to elephant trunk guidance (23,24), and beak-manipulated tool use in birds (14). They reveal lateralization as a common strategy for visual guidance. Despite these similar strategies, the causal dynamics between the eye and appendage differ across species (25,26), developmental history (2,27–30), and task geometry (11,31,32). In addition, perturbation experiments such as monocular occlusion (33) demonstrate varying plasticity in the underlying control axis (10,22,34). Expanding the comparative range to include invertebrates (3,35–37) thus offers an ideal opportunity to study whether lateralized control axes represent a conserved solution across very diverse taxa, and more specifically, how the size-limited nervous systems of insects establishes causality and implements plasticity in similar visual appendage-guidance tasks.

Few insect species display continuous visual guidance of appendages (38–41), similar to reaching and grasping behaviors studied in vertebrates. Consequently, most investigations of lateralization in insects focused on discrete quantifications of actions, such as turning direction and leg use (42). One emerging model for continuous eye-appendage coordination is the diurnal hummingbird hawkmoth (*Macroglossum stellatarum*), which uses visual information to control both body and proboscis movements (43,44), guiding their proboscis to flower nectaries while hovering (45–48). Their central appendage, under bilateral visual control, mirrors the control system of the bird beak (14,20,49) and the elephant trunk (23,24,50), providing a unique analogous model to unpaired appendage control in vertebrates.

We here used the flower probing behaviour of the hummingbird hawkmoth to investigate the mechanisms of sensory-motor lateralization in the continuous control of the proboscis appendage. Combining high-speed videography, markerless pose estimation, and gaze estimation revealed the stability of proboscis lateralization, its functional consequences, and a time-resolved relationship between the eye and the proboscis. Additionally, we performed monocular occlusion to investigate the plasticity of the hawkmoth visuo-motor axis and the causal dynamics of its components.

## Results

### Probing hawkmoths displayed a continuous range of proboscis lateralization

Using markerless pose estimation on high-speed recordings, we analysed hawkmoths probing on artificial flowers with a stripe pattern (Fig. 1A). We quantified the relative position of the proboscis tip to the head, in head axis coordinates, revealing whether hawkmoths kept their proboscis centred, or left or right of the midline during probing (Fig. 1*C-D*). At a group level (n=95, N=4530; total contacts=610,450), the distribution of all proboscis positions was symmetric in the lateral direction (Fig. 1*E*), and did not differ significantly from 0, the centre of the head axis (Fig. 1*F*; Wilcoxon signed-rank compared to zero, p>0.05). However, analyzing the distributions of individual animals revealed distinct lateralization subgroups, with animals whose proboscis positions were shifted to the left or right of the head midline (Fig. 1*I*). To analyse whether these shifts were consistent, we compared the median proboscis positions of all probing bouts (see Fig. 1*B*) an individual performed within an experimental session to zero (Wilcoxon signed-rank test, unbiased p>0.05, lateralized p<0.05). Animals with significant lateralization were separated into left- (n=32 moths, N=1626 bouts) and right-lateralized (n=25 moths, N=1391 bouts) subgroups (‘left’ and ‘right’ group from here on), revealing respective shifts in the subgroup proboscis position distributions (Fig. 1*G-H*; Wilcoxon signed-rank compared to zero, two-tailed, p<0.001; between-group: Wilcoxon rank-sum, p<0.001). All animals, including those with significant lateralization, displayed a considerable variation in bout-wise proboscis positions, which in many animals included individual bouts with distribution maximums in the opposite direction of their median lateralization tendency. Thus, many significantly right-lateralized hawkmoths also had left-shifted proboscis distributions in individual probing bouts, and vice versa. The resulting distribution of per-animal lateralization depicted a continuous range of lateralization amplitude in world units (millimeters), with similar proportions of lateralization subgroups (Fig. 1*I*; left n=32, right n=25, unbiased n=38). Measuring relative proboscis position with a lateralization index (see *Methods*) further demonstrated animal-specific proboscis lateralization and a similar continuous range of lateralization strengths (Fig. 1*J*). We confirmed that these distributions were not the result of sampling bias by randomly resampling the bout position maximums within and across animals, and found that the resampled distributions significantly differed from the data (see *Methods*, Supplementary Fig. S1*D*–*H*).

**Figure 1:**
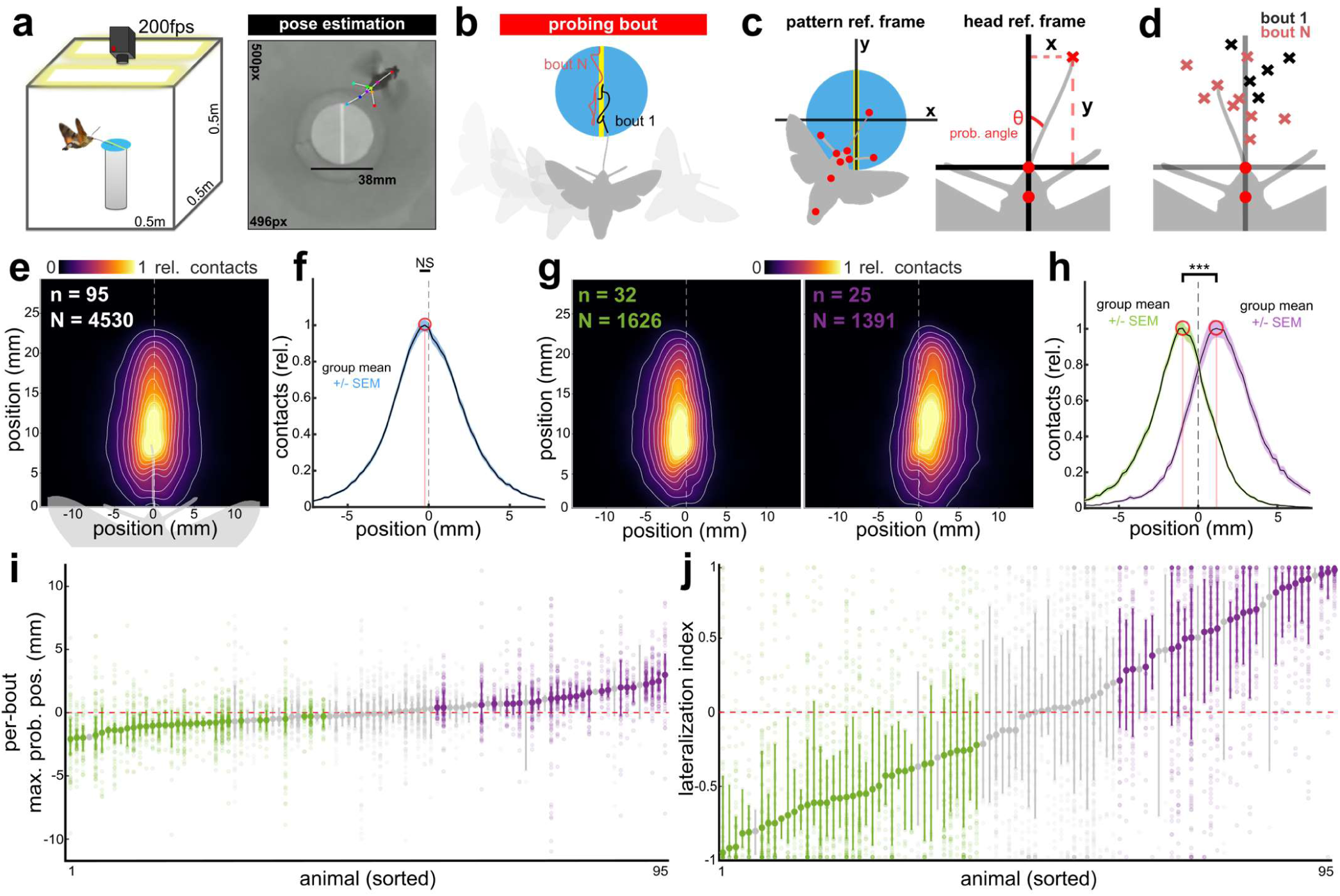
Quantification of lateralization in the hunningbird Hawkmoth proboscis. **A**: Schematic of behavioural experiment setup (left). Example labelled frame from recorded videos with DeepLabCut-predicted keypoints (right). **B:** Schematic depicting the course of a probing bout. Coloured paths indicate proboscis trajectories from different probing bouts. **C:** Schematics depicting the two primary reference frames used in this study. The pattern reference frame (left) has the origin on the centre of the stimulus pattern, while the head reference frame (right) has the origin on the base of the proboscis. **D:** Example proboscis positions from two distinct probing bouts, using the head reference frame. Colours depict proboscis positions from separate bouts. **E:** Positions of the proboscis relative to the proboscis base during stimulus contact, normalized and averaged across individuals (n) and bouts (N). **F:** Normalized distribution of proboscis positions along the x-axis, indicating leftwards (negative) and rightwards (positive) proboscis positions. Distribution maximum circled in red for visualization. Distribution of per-animal medians was compared to zero using the Wilcoxon signed-rank test. **G-H:** Proboscis position heatmaps (G) and lateral position distributions (H) of significantly left- (green; left) and right-lateralized (purple; right) hawkmoths. Distributions of per-animal medians were compared to one another with the Wilcoxon rank-sum test. **I-J:** Distributions of per-bout proboscis amplitudes (I) and per-bout lateralization strengths (J), for each animal. Large opaque circles indicate distribution median, with vertical lines depicting 95% CI. Small translucent circles are per-bout distribution maximums (I) or lateralization indices (J). Each animal’s distribution was compared independently to zero using Wilcoxon signed-rank test. Colours depict lateralization significance and direction (green=significantly leftwards, purple=significantly rightwards, gray=no significance/unbiased). Statistical test results are abbreviated as * (p<0.05), ** (p<0.01), *** (p<0.001), and NS (not significant), with full details available in Tables S8-S12.

### Proboscis lateralization was present at eclosion and consistent across days

To determine whether the observed proboscis lateralization was consistent across days, we analysed the median bout positions of feeding-experienced animals, which had fed from gravity feeders but not the artificial flowers before, across 3 to 5 consecutive days (first three days shown for visualization in Fig. 2*A* – upper half; left n=19, right n=11, unbiased n=12; all days shown in Supplementary Fig. S2*A*).

**Figure 2:**
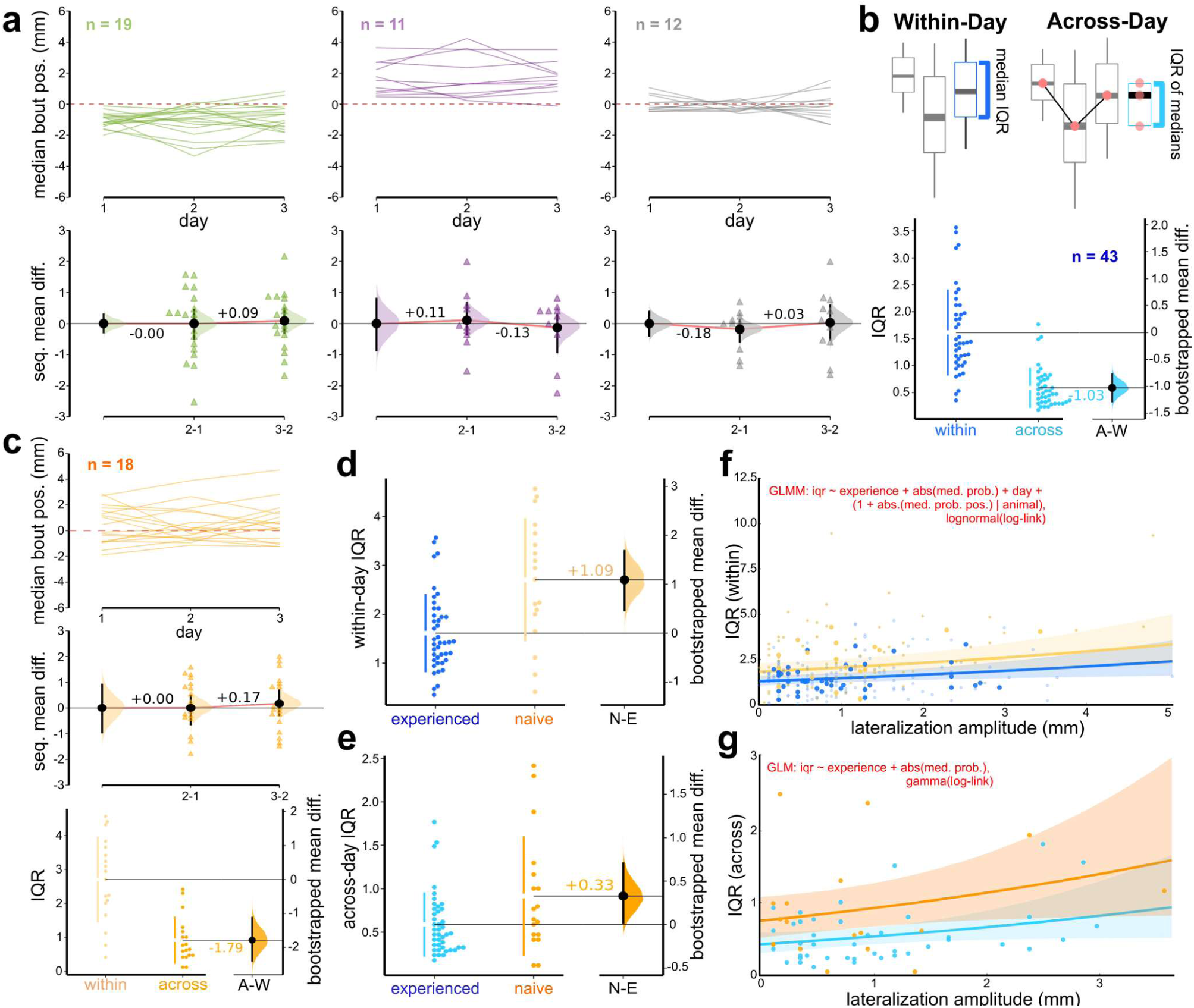
Proboscis lateralization is a consistent and innate feature. **A**: (top) Sequential medians of proboscis lateralization amplitude across days. Each line represents an individual hawkmoth’s median lateralization amplitude. Colours indicate lateralization subgroup. (bottom) Sequential mean difference of the group mean amplitude per day. Triangles represent individual animals’ differences in amplitude between consecutive days. Black circle and vertical bars represent mean difference and 95% CI. Shaded distribution depicts bootstrapped mean difference reference distribution. **B:** Comparison of variability in feeding-experienced hawkmoths through two measures of interquartile range (IQR). Circles represent within- and across-day IQR of each animal. Vertical lines depict IQR of the distribution. Group means depicted by horizontal lines. Bootstrapped distribution depicts effect size of difference in means between both measures. **C:** Consistency and variability of naïve hawkmoths. **D-E:** (D) Bootstrapped mean difference of within-day IQR between experienced (blue) and naïve (orange) hawkmoths. (E) Bootstrapped mean difference of across-day IQR between experienced and naïve hawkmoths. **F:** Generalized linear mixed model predicted fit to the within-day IQR of all animals, as a function of lateralization amplitude. Lines depict back-transformed predicted fit of each hawkmoth group (yellow=naïve, blue=experienced). Shaded bands depict 95% CI of fit. Large opaque points depict overall median within-day lateralization amplitude; small points depict median amplitude per day. **G:** Generalized linear model predicted fit to the across-day IQR of all animals, as a function of lateralization amplitude. Lines depict back- transformed predicted fit of each hawkmoth group (orange=naïve, cyan=experienced). Shaded bands depict 95% CI of fit. Points depict overall median lateralization amplitude across days, per animal.

Feeding-experienced hawkmoths overwhelmingly retained the proboscis lateralization direction displayed on the first day of testing, across multiple days (Fig. 2*A*; Supplementary Fig. S2*C*). To assess whether the proboscis lateralisation amplitude remained consistent across days, we compared the variation in the median per-bout proboscis positions within days, to the variation in median position across-days (Fig. 2*A* – lower half). Since proboscis positions varied significantly more within a day than across days (see *Methods*, Fig. 2*B*; -1.79 [95% CI: -2.4, -1.12]), we concluded that proboscis lateralization was consistent across days in feeding-experienced hawkmoths.

To investigate whether the proboscis lateralization of experienced animals was innate, or potentially acquired while using the proboscis to feed from gravity feeders, we quantified the proboscis lateralization of freshly-eclosed hawkmoths which were fully naïve to any feeding. Similar to experienced animals, naïve hawkmoths displayed significant lateralization on the first day of testing (Fig. 2*C*; n=18; Supplementary Fig. S2*A*,*C*). Within-day variability of the median proboscis position was also significantly greater than across-day variability, indicating that consistent lateralization was present from eclosion. Comparing variance measures between the two groups revealed that naïve animals had significantly higher variability than experienced ones within a day (1.09 [95% CI: 0.47, 1.68]), albeit less strongly across days (0.328 [95% CI: 0.0503, 0.705]) (Fig. 2*D*-*E*). Thus, while naïve hawkmoths displayed significant innate proboscis lateralization, their amplitudes were more variable compared to experienced hawkmoths.

We next investigated whether lateralization amplitude impacted the variability of proboscis positioning by fitting mixed models to the within- and across- day variance measures (see *Methods*). Increasing lateralization amplitude – measured as the daily absolute median of bout maximums – resulted in significantly increased estimates of within-day variability for both groups (Fig. 2*F*; Table S1A-B). Thus, animals with more lateral median proboscis positions had a higher variability in these positions across probing bouts. Across days, lateralization amplitude did not significantly impact estimates of variability (Fig. 2*G*; Table S2B). Testing day as a factor did not have a significant impact on variability, but predicted significantly increased variability on the third day of testing (Table S1A). Conversely, we found that increasing lateralization strength (1 being the maximum absolute strength) resulted in a reduction of variability, on a within-day, but not across-day level (Supplementary Fig. S2*D-E*; Table S3-S4).

Together, these results demonstrate that while proboscis lateralization might have an experience component, suggested by the increased variability of naïve animals over experienced ones, it was present on the first day and consistent across days in both naïve and experienced hawkmoths.

### Proboscis lateralization was linked to feature probing strategies

Given that hawkmoths displayed innate and consistent proboscis lateralization, we next asked: what are its functional consequences for flower probing? We therefore analysed the proboscis contacts of the inspecting hawkmoths on the flower, in particular on the stripe pattern, which they were targeting (see right-lateralized hawkmoths in Fig. 3*A*).

**Figure 3:**
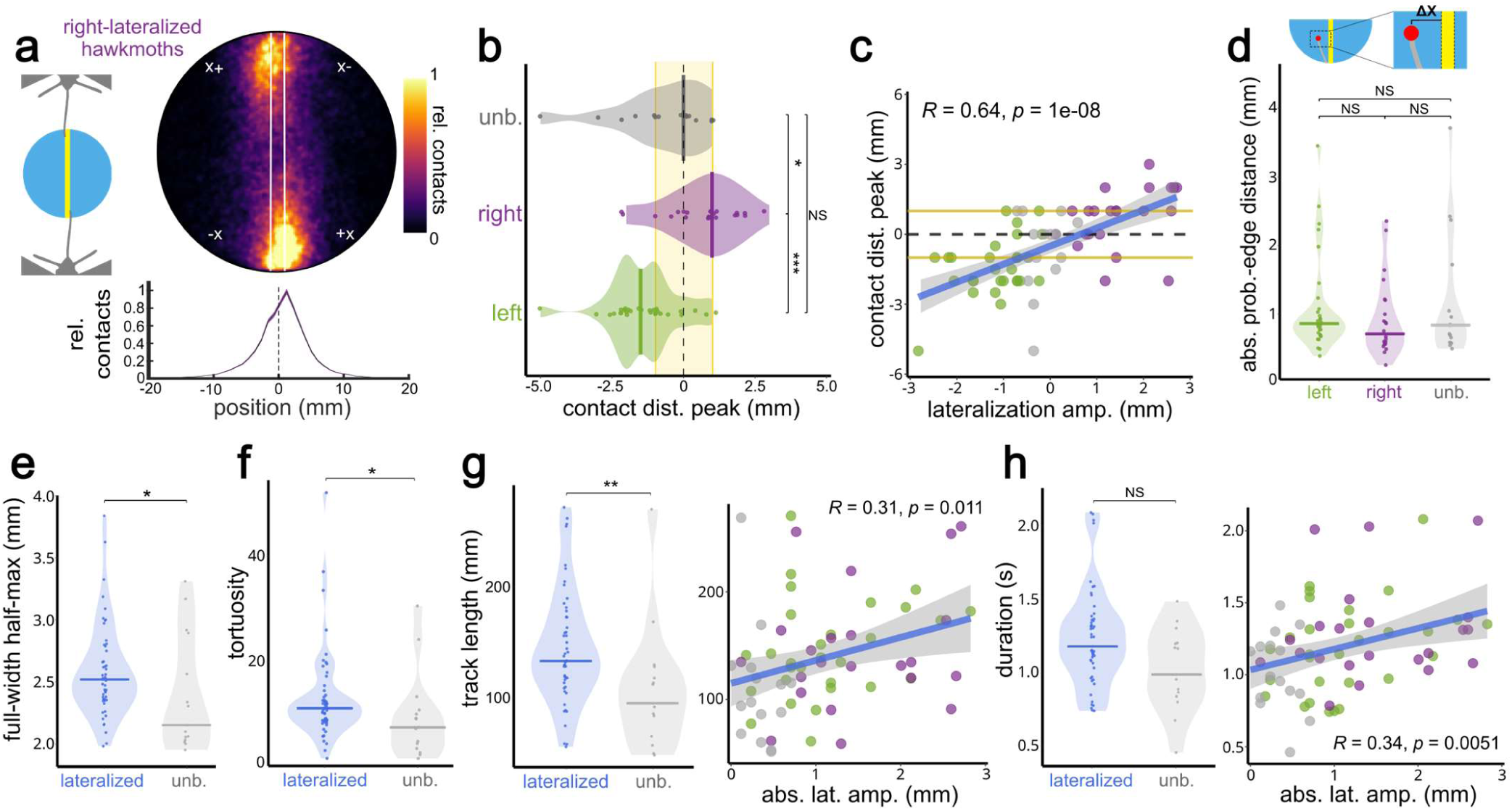
Lateralization is coupled to feature targeting and trajectory characteristics. **A**: Example of hawkmoth proboscis contact positions using right-lateralized hawkmoths. (left) Schematic depicting feature targeting of the rightwards edge. (top) Normalized and averaged proboscis contact positions on the stimulus surface. White lines depict stripe edges. (bottom) Normalized distribution of contact positions relative to the stripe midline (midline=0mm; edges=+/-1mm). **B:** Per-animal medians of the contact distribution maximum across bouts. Points indicate median bout- wise contact position per animal. Group colours depict lateralization direction. Short vertical bars depict group-level median. Shaded violin regions depict density of group measurements. Shaded yellow region depicts the position of the stripe on the stimulus. Statistical test results indicate corrected pairwise Wilcoxon tests. **C:** Per-animal median of the contact distribution maximum across bouts, as a function of lateralization amplitude (mm). Points depict animals coloured by subgroup. Yellow lines depict position of the pattern stripe. Black dashed line depicts the pattern midline. Blue line and shaded region depict fitted linear regression and confidence intervals. **D:** Accuracy of contact positions. Points indicate per- animal medians of the absolute distance between the proboscis contact position and the nearest stripe edge. Statistical test results indicate corrected pairwise Wilcoxon tests. **E-F:** Comparisons between lateralized and unbiased hawkmoths for (E) precision as full width at half maximum of contact distributions (mm), and (F) proboscis trajectory tortuosity. Points represent bout-wise median of respective measure per animal. Colour depicts subgroup type (blue=lateralized, gray=unbiased). Distributions within each measure were compared to one another using the Wilcoxon rank-sum test. **G:** (left) Probing trajectory track length (mm), points represent bout-wise median per animal; (right) track length as a function of absolute lateralization amplitude (mm), fitted with a linear regression. **H:** (left) Duration of probing bout (sec), points represent bout-wise median per animal; (right) bout duration as a function of absolute lateralization amplitude (mm), fitted with a linear regression. Statistical test results are abbreviated as * (p<0.05), ** (p<0.01), *** (p<0.001), and NS (not significant), with full details available in Tables S8-S12.

Interestingly, we found that lateralized hawkmoths aimed their proboscis such that it contacted near the stripe edge (e.g. -1/+1 mm of pattern midline), preferentially on the side corresponding to their proboscis lateralization direction (Fig. 3*A*-*B*; left n=29, right n=22, unbiased n=15; Kruskal-Wallis test, p<0.01; corrected pairwise Wilcoxon rank-sum, p<0.05). Unbiased hawkmoths switched between contacting either edge, resulting in more centered bout-wise distributions (Fig. 3*B*). The number of bouts did not significantly differ between groups, indicating an equal propensity to interact with the stimulus (Supplementary Fig. S3*C*; Kruskal-Wallis test, p>0.05).

Feature targeting was found on the first day of testing for all three groups, which confirmed the early presence of visual targeting strategies (Supplementary Fig. S3*A*). Peak proboscis contact positions of the experienced hawkmoths linearly transitioned with lateralization amplitude, reinforcing that feature targeting was significantly linked to lateralization (Fig. 3*C*; r=0.64, p<0.001).

As the stripe edges were a strongly relevant feature for the hawkmoths, we quantified how accurate proboscis placements were relative to the nearest edge. None of the three groups had significant differences in median bout-wise edge distance (Fig. 3*D*; Kruskal-Wallis test, p>0.05), indicating equal accuracy with proboscis targeting. Given that accuracy did not differ between lateralized subgroups, we pooled left and right hawkmoths (n=51) to investigate whether being lateralized resulted in greater precision with proboscis placement. Comparing precision, quantified as full width at half maximum of the contact distribution (mm), between the lateralized and unbiased groups demonstrated that unbiased hawkmoths were more precise than lateralized hawkmoths (Fig. 3*D*; Wilcoxon rank-sum, p<0.05). This precision was not affected by lateralization amplitude (Supplementary Fig. S3*B*; r=0.17, p=0.18).

Having found that precision, but not accuracy, was impacted by lateralization, we next investigated if lateralization affected parameters of the probing trajectory. At a group level, we found that lateralized hawkmoths produced more tortuous proboscis trajectories (Fig. 3*F*; Wilcoxon rank-sum, p<0.05), but this tortuosity did not smoothly increase with lateralization amplitude (Supplementary Fig. S3*E*; r=0.13, p=0.3). Consequently, we also found that lateralized hawkmoths produced longer trajectories than unbiased hawkmoths, which did linearly increase with absolute lateralization amplitude (Fig. 3*G*; Wilcoxon rank-sum, p<0.01; r=0.31, p=0.011). Despite these longer trajectories, probing duration was not affected by lateralization subgroup, but did linearly increase with lateralization amplitude (Fig. 3*H*; Wilcoxon rank-sum, p>0.05; r=0.34, p=0.0051). We also found that probing speed of the proboscis did not differ between groups either categorically or continuously (Supplementary Fig. S3*D*,*F*; Wilcoxon rank-sum, p>0.05; r=0.15, p=0.22), indicating that probing characteristics were not simply a result of motor impairment.

Taken together, we found a clear differentiation in feature probing strategy between lateralized and unbiased hawkmoths, with unbiased hawkmoths showing higher precision in probing, while lateralized ones had longer and more tortuous trajectories.

### Hawkmoths viewing strategies are coupled to proboscis lateralization

Since feature probing, for which we found diverging strategies between lateralized and unbiased hawkmoths, is a visual control task (43), we hypothesized that lateralization may be driven by stimulus viewing strategies. To assess the viewing strategies, we first analysed the hawkmoths’ head positions relative to the stripe. The relative occupancy of the tracked head keypoint was highest on either end of the vertical axis of the stripe (Fig. 4*A*-*B*; n=45, N=358,320 frames). Qualitatively, difference heatmaps revealed left and right hawkmoths positioned themselves on opposite sides of the stripe (Fig. 4*C*; Supplementary Fig. S4*A*; left: n=17, N=152,099; right: n=11, N=105,451), which was supported by robust mixed model analysis (Fig. 4*D*; Table S3; see *Methods*): in left- lateralized moths, both the proboscis base and head were displaced significantly rightwards relative to the proboscis tip (Table S5C), and each lay right of the stripe midline (Table S5B). In contrast, right- lateralized moths showed the reverse: the base and head trended to be shifted left of the midline (Table S3B), and were significantly left of the proboscis (Table S5C). Unbiased moths exhibited no reliable offsets (all 95% CIs widely overlapping zero), indicating alignment with the midline, and alignment between body parts (Table S5B-C). Between left- and right-lateralized groups, these displacements differed sharply: compared to left moths, the proboscis base and head were significantly more left-shifted in the right group, whereas the average proboscis tip estimate trended to be more rightwards (Table S3D). The angle of the head (Supplementary Fig. S4*B*-*D*), and that between the proboscis base and the pattern center (Supplementary Fig. S4*E*-*G*), did not differ between groups. The joint head-pattern angle was centred on the pattern (Supplementary Fig. S4*E*- *G*), indicating that all hawkmoths probed the pattern with similar ‘attack’ angles, but were laterally shifted off the midline in head and body position corresponding with their lateralization group.

**Figure 4:**
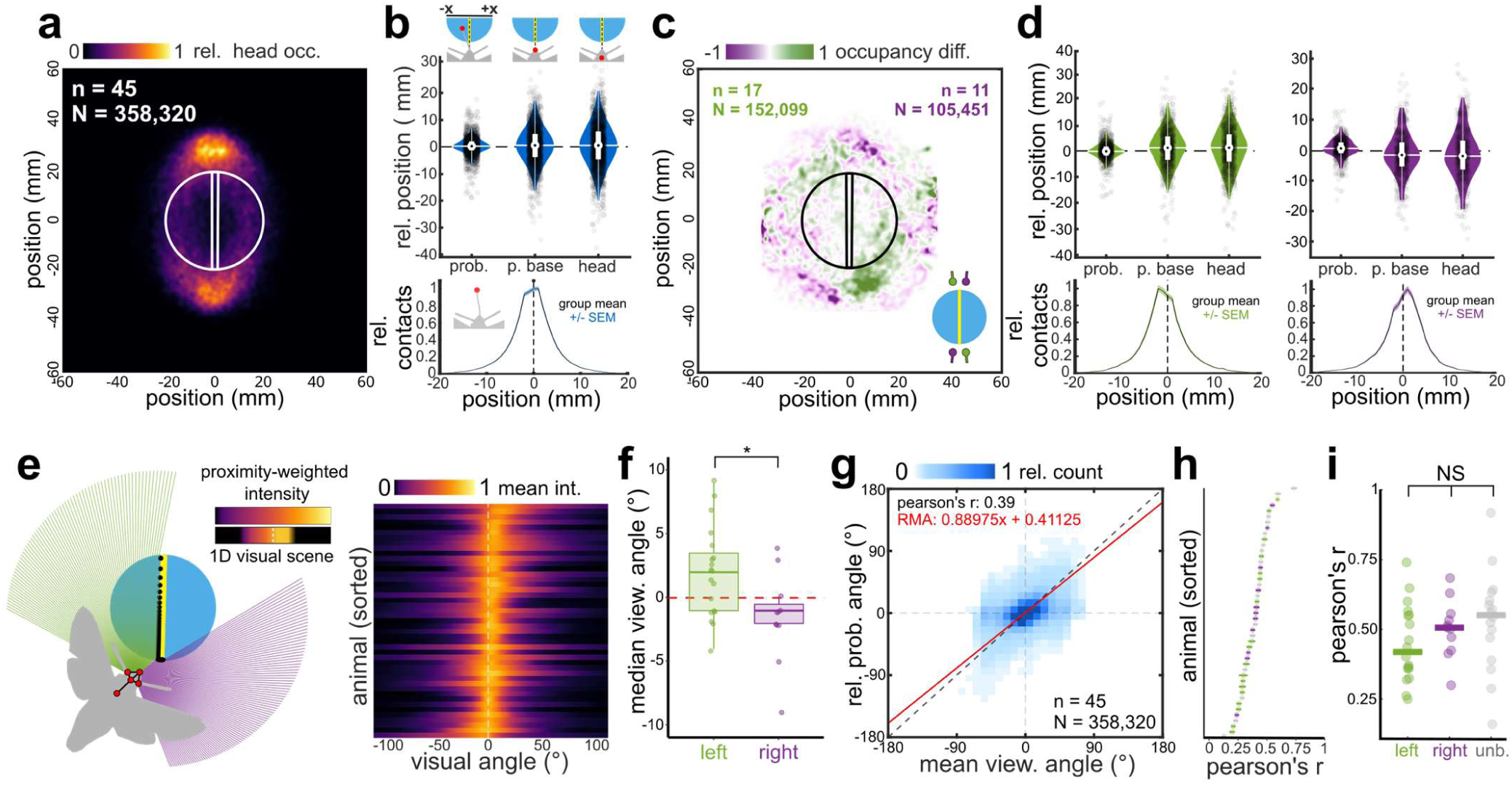
Position and gaze estination of probing Hawkmoths revealed a lateralized eye-proboscis axis. **A**: Normalized and averaged occupancy of the tracked head keypoints across all moths (n) and frames (N) where the proboscis was in contact. White outline depicts the stimulus location. **B:** (top) Per-bout average positions (mm) of three tracked keypoints: proboscis top, proboscis base, and head, relative to the stripe midline. Points depict bout averages, white boxes depict IQR about the group mean (innermost black point). (bottom) Proboscis contact position distribution relative to stripe midline for all contacts. **C:** Normalized and averaged difference heatmap depicting the difference in head occupancy between left-lateralized (green; n=17, N=152,099) and right-lateralized (purple; n=11, N=105,451) hawkmoths. Inset schematic depicts which group more dominantly occupies the frame quadrant. **D:** Per-bout average positions of keypoints for left-lateralized (left) and right-lateralized (right) hawkmoths. **E:** (left) Schematic depicting the visual field used for the gaze estimation of probing hawkmoths (see *Methods*). (right) Averaged visual scene intensities for experienced hawkmoths, sorted in ascending order based on angle of maximum intensity. **F:** Distributions of median bout-wise viewing angle (deg), per animal. Points depict individual animals’ medians, separated into left- and right-lateralized subgroups. Statistical comparison performed using Wilcoxon rank-sum test. **G:** Density scatterplot of per-contact mean viewing angle and relative proboscis angles across all moths (n) and contacts (N). Red line depicts fitted Reduced Major Axis regression. Dotted black line depicts 1:1 relationship. **H:** Pearson correlation strengths of per-contact mean viewing angle and relative proboscis angle, per animal. Animals sorted in order of ascending correlation strength. Colours of points depict lateralization direction. Horizontal bars within points depict 95% CI of correlation coefficient. **I:** Pearson correlation strengths separated into lateralization subgroups. Horizontal bars depict group median. Groups were compared using the Kruskal-Wallis test. Statistical test results are abbreviated as * (p<0.05), ** (p<0.01), *** (p<0.001), and NS (not significant), with full details available in Tables S8-S12.

Such positioning could have resulted in one eye receiving relatively more visual input, allowing hawkmoths to both view the pattern and guide the proboscis preferentially in one visual axis. To quantitatively assess this hypothesis, we simulated the visual field of each eye during all proboscis contacts and calculated the proximity-weighted intensity of the pattern stimulus at each visual angle (Fig. 4*E*), relative to their head coordinate system (see *Methods*). This resulted in a per-frame mean viewing angle (the weighted mean angle indicating the relative position of the pattern to the moth’s eyes; see *Methods*), as well as an overall visual scene intensity (i.e. at each angle of the visual field, what is the relative visual signal strength at any given time; see *Methods*). The weak relationship between head angle and mean pattern viewing angle (Supplementary Fig. S4*K*; r=0.3, p=0.049) suggested that our gaze estimation approach captured different visual information than what solely using head orientation would have estimated. The relative intensity across visual angles differed between moths, and predominantly consisted of maximum intensities in the frontal zone (-10°:10°; V-test against 0°, p<0.001) (Fig. 4*E*-*F*). Consistent with our hypothesis, bout-wise analysis revealed that both the left and right groups were significantly shifted from one another in the angles at which these maximum intensities occurred (Fig. 4*F*; Wilcoxon rank-sum, p<0.05). We also found a significant correlation between the overall angle of maximum intensity, and the median proboscis angle, at a per-animal level (Pearson r=0.36, p=0.016; Fig. S4*L*). This revealed an individualised relationship between lateralized visual input and lateralized proboscis positioning, linking the eye to the proboscis.

To investigate this relationship further, we analysed the per-frame mean viewing angle and relative proboscis angle for each animal (see *Methods*). At a group level (n=45, N=358,320 contacts), we found a significant relationship between viewing angle and proboscis angle (r=0.39, p<0.001) (Fig. 4*G*). Fitting a reduced major axis regression further indicated a structural, albeit noisy, relationship between the two instantaneous angles (RMA=0.89x + 0.41) (Fig. 4*G*). Per-animal, we found a wide distribution of significant Pearson correlations ranging from 0.16 to 0.8 (Fig. 4*H*). Specific lateralization direction did not affect the relationship strength between proboscis angle and visual input (Kruskal-Wallis test, p>0.05) (Fig. 4*H*-*I*).

In summary, we found that hawkmoths positioned themselves relative to the visual pattern according to their proboscis lateralization, and had a time-locked relationship between their proboscis angle relative to the head, and the visual angle at which the pattern was viewed, forming a lateralized eye- proboscis axis.

### Hawkmoths flexibly adjusted viewing strategies, but not proboscis lateralization, upon monocular occlusion

Having established that lateralized hawkmoths show a preferred eye-proboscis axis, we sought to determine its mechanistic basis through monocular occlusion of the eyes. More specifically, does the eye drive the proboscis, or does the proboscis drive the eye?

To investigate this, we painted the frontoventral region of the eye corresponding to the moths’ proboscis lateralization direction, to interrupt the eye-proboscis axis on their preferred side (Supplementary Fig. S5*A*). We found that hawkmoths generally did not shift their proboscis placement into view of the non-occluded eye (Fig. 5*A*; left eye painted n=12, right eye painted n=9). Although some individuals shifted their proboscis placement more towards the direction of the free eye (Supplementary Fig. S5*B*, Wilcoxon signed-rank, p<0.05), the effect size indicated there were no significant differences after painting for either group (Fig. 5*B*; paired Hedges’ g, left: +0.09mm [- 0.28,0.45], right: -0.57mm [-1.25,0.42]). Surprisingly, some individuals shifted their proboscis even more strongly towards their initial lateralized direction (Fig. 5*B*; Supplementary Fig. S5*B*, Wilcoxon signed-rank, p<0.05), potentially seeking to place the proboscis in the unpainted lateral region of the occluded eye. Painted hawkmoths still targeted the initially preferred stripe edge, further indicating that the proboscis strategy remained consistent after painting (Supplementary Fig. S5*D*).

**Figure 5:**
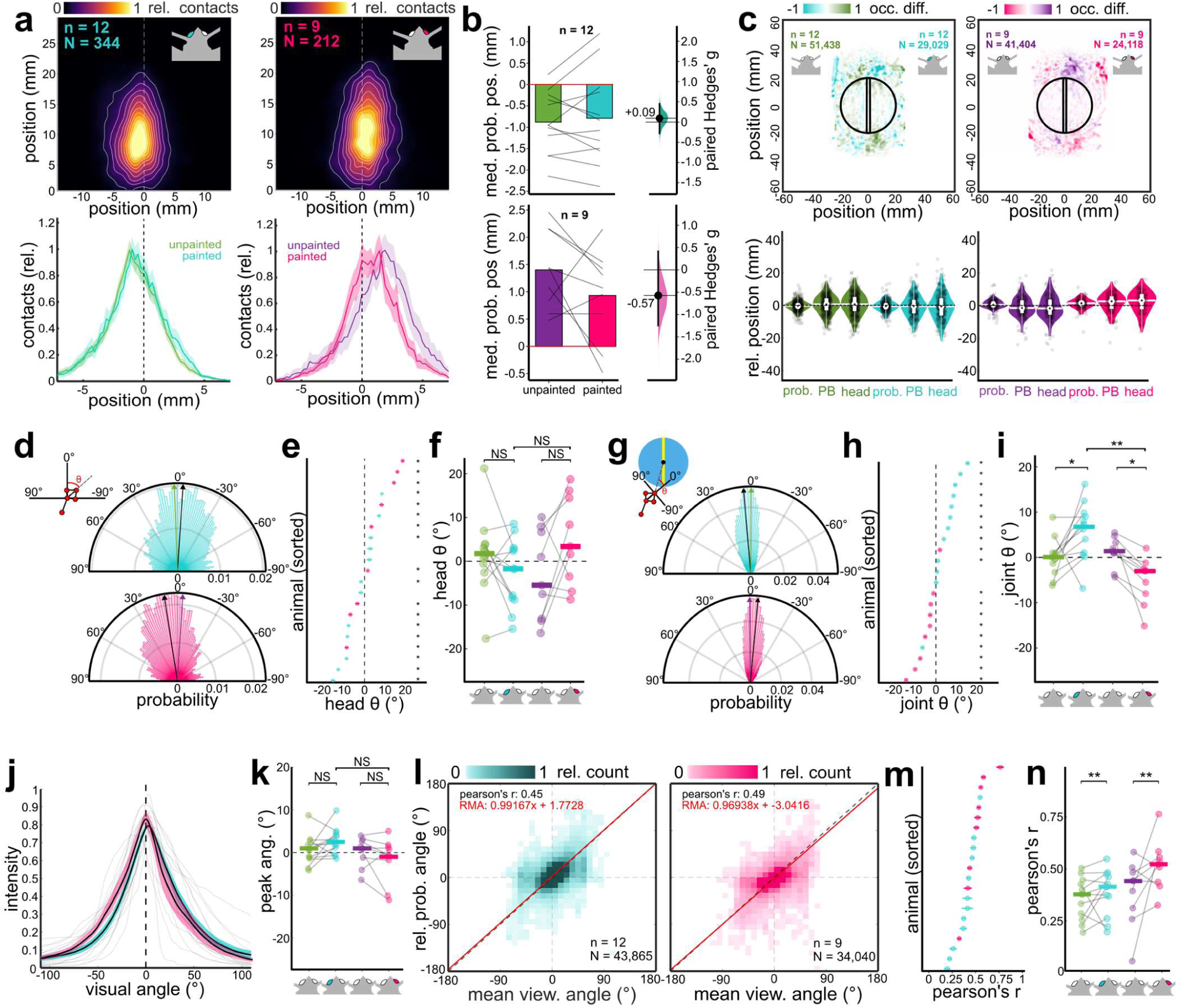
Monocular occlusion shifts Hawkmoth positioning and strengthens the lateralized eye-proboscis axis. **A**: (top) Proboscis positions relative to proboscis base of monocularly-occluded hawkmoths (left: left eye painted, teal; right: right eye painted, magenta), normalized and averaged across individuals (n) and bouts (N). (bottom) Normalized distribution of proboscis positions along the x-axis (mm), with before- and after-occlusion distributions depicted by colour. **B:** Median proboscis lateralization amplitudes (mm) before and after eye painting, for all moths within subgroups (n). Lines depict per-animal differences. Filled bars depict sign and magnitude of group mean. Bootstrapped paired Hedges’ g depicts effect size of difference in mean after painting. **C:** (top) Normalized and averaged difference heatmaps depicting the difference in head occupancy before and after painting for both lateralized subgroups (left: left-lateralized before and after, n=12, N=51,438; n=12, N=29,029; right: right-lateralized before and after, n=9, N=41,404, n=9, N=24,118). (bottom) Bout- wise average positions of tracked keypoints (mm) before and after painting, for both subgroups. **D:** Distributions of head orientation during all proboscis contacts following occlusion. Coloured bars depict histogram probability. Black arrows depict painted group median angle. Coloured arrows depict group median angle before painting. **E:** Per-animal mean head orientation, sorted in ascending order. Points are coloured by eye painted. Horizontal bars within points depict 95% CI of mean angle. Individual means tested for significance using the one-sample circular mean test. **F:** Mean head orientation angle before and after painting, for both lateralization subgroups. Horizontal bars depict group medians. Within-group comparisons were made using Wilcoxon signed-rank, between-group comparisons were performed with Wilcoxon rank- sum. **G:** Distributions of joint angles between proboscis base and pattern centre. **H:** Per-animal joint angle, sorted in ascending order. **I:** Mean joint angle before and after painting. Within-group comparisons were made using Wilcoxon signed- rank, between-group comparisons were performed with Wilcoxon rank-sum. **J:** Averaged visual scene intensity distributions. Coloured confidence intervals indicate painted group. **K:** Comparison of maximum intensity angles before and after painting. Horizontal bars depict group medians. Within-group comparisons were made using Wilcoxon signed- rank, between-group comparisons were performed with Wilcoxon rank-sum. **L:** Density scatterplots of per-contact mean viewing angle and relative proboscis angles across all moths (n) and contacts (N) for the left and right painted groups, respectively. **M:** Pearson correlation strengths of per-contact mean viewing angle and relative proboscis angle, per animal. Animals sorted in order of ascending correlation strength. Colours of points depict painted group. **N:** Pearson correlation strengths before and after monocular occlusion. Horizontal bars depict group median. Within-group comparisons were made using the Wilcoxon signed-rank test. Statistical test results are abbreviated as * (p<0.05), ** (p<0.01), *** (p<0.001), and NS (not significant), with full details available in Tables S8-S12.

Hawkmoths strongly re-positioned themselves depending on which eye was painted (Fig. 5*C*; left- unpainted: n=12, N=51,438; left-painted: n=12, N=29,029; right-unpainted: n=9, N=41,404; right- painted: n=9, N=24,118). Similar to the unpainted moths (Fig. 4*J*), robust mixed model estimates revealed mirrored lateral displacements across groups: In left-unpainted (LU) moths, the proboscis base and head tend to lay right of the midline (Table S4B) and were significantly rightward relative to the proboscis (Table S6C). After painting (LP), these offsets disappeared (Table S6B-C), indicating a leftward shift in body position. Right-unpainted (RU) moths showed the opposite configuration, with the base and head (Table S6B) tending to shift leftward of the midline, and significantly left of the proboscis (Table S6C). In right-painted (RP) moths, these leftward displacements weakened (Table S6B-C), indicating a rightward shift in body position. Between-group differences (LP–LU, RP–RU) were directionally consistent but not statistically significant (Table S6D). Thus, occluding the eye ipsilateral to the proboscis lateralization direction caused hawkmoths to shift position in the direction of the non-occluded eye, thus indicating a potential change in viewing strategy.

While the head orientation angles (Fig. 5*D*) did not differ significantly between painted groups (Fig. 5*F*, Wilcoxon signed-rank, p>0.05; Wilcoxon rank sum, p>0.05), the proboscis base - pattern angles (Fig. 5*I*) of both lateralized groups, at an individual and group level, showed significant differences between conditions, demonstrating that the centre (and thus the vertical axis) of the stripe was shifted to be more in view of the painted eye (Fig. 5*I*, Wilcoxon signed-rank, p<0.05; Wilcoxon rank-sum, p<0.01). Correspondingly, the correlation between the angle of maximum scene intensity and the median proboscis angle strengthened after occlusion (Fig. S5*C*; r=0.47, p=0.033). The distribution of average visual scene intensities and peak angles did shift as well, although not statistically significantly (Fig. 5*J*-*K*).

At a per-frame level, readjustments of the head position drove a stronger coupling between mean viewing angle and relative proboscis angle (Fig. 5*L*, compared to that seen in Fig. 4*H*), increasing the overall correlation significantly for both lateralized groups (Fig. 5*N*, Wilcoxon signed-rank, p<0.01). Intriguingly, painting the non-preferred eye did not induce significant changes in body positioning nor proboscis lateralization (Supplementary Fig. S5*E-F*,*H-I*; Wilcoxon signed-rank p>0.05; Wilcoxon rank sum p>0.05), but did result in noticeable orientation changes (Supplementary Fig. S5*O*; Wilcoxon signed-rank p<0.01; Wilcoxon rank sum p<0.05). Despite these changes, peak viewing angles did not significantly shift between conditions (Supplementary Fig. S5*P*-*Q*), nor did the correlation between mean viewing angle and proboscis angle (Supplementary Fig. S5*R-T* compared to Fig. 4*H*).

Monocular occlusion thus revealed that hawkmoths flexibly reconfigured their viewing posture around a fixed proboscis motor strategy, which maintained a preferred eye-proboscis axis. Occluding the contralateral – or ‘non-preferred’ – eye resulted in adjustments to head orientation, but none of the components of the proboscis-eye axis.

## Discussion

In this study, we demonstrated that hummingbird hawkmoths exhibit stable, individual-level lateralization during visually guided control of the proboscis, revealing lateralized control of an unpaired appendage in an invertebrate. Unlike most studies of insect lateralization, which focus on discrete actions such as turning or limb selection, our results describe continuous sensory-motor coordination, allowing the dynamics of eye-appendage coupling to be quantified over time. This establishes hawkmoths as a tractable system for studying lateralized visuo-motor control in a continuous behavioural task.

### Lateralization of sensory input and visuo-motor coupling

The individual-level proboscis lateralization we found in the hummingbird hawkmoth, spanning different directions, strengths, and amplitudes (Fig. 1I-J), demonstrates the first evidence of lateralized unpaired appendage control in an invertebrate. Hawkmoths thus provide a unique comparative model for the lateralized control of unpaired appendages in vertebrate taxa, such as elephant trunk use (23,24) and beak-guided tool manipulation in birds (14). Despite their vast evolutionary distance, all three animal models share a control problem which is different to that of bilaterally paired appendages: how does an organism guide a centrally positioned effector using input from paired sensors?

Time-resolved analysis revealed that proboscis placement and pattern viewing angle were significantly correlated, indicating that hawkmoths maintained a preferred visuo-motor axis linking the eye, the appendage, and the visual target (Fig. 4G-I; Supplementary Fig. S4L). Adopting a dominant eye to guide asymmetric motor control may provide one solution to the control problem: preferential neural pathways linking one eye to the appendage could reduce the routing required for control, in both computation and anatomy (16,19,21). Unbiased hawkmoths demonstrated similar instantaneous coupling between viewing angle and proboscis position, placing both the target and appendage in the same visual field, but switching frequently between sides (Fig. 4I). Accordingly, this suggests that also hawkmoths lacking consistent proboscis biases establish temporary visuo-motor axes during probing, which guide proboscis movements within an interaction, but flexibly adjust over consecutive bouts. Similar coordination has been shown in humans and elephants, where the eye forming the axis is flexibly shifted based on appendage geometry and motor task (11,23,32).

In other insects, lateralized visual behaviours have been described for tasks such as gap crossing (51,52), turning (53–57), and escape responses (58,59), but consistent lateralized coupling between eye use and limb or body movements has not been demonstrated. In locusts, for example, population-level eye preference during predator surveillance does not reliably predict escape direction (58), indicating that sensory lateralization alone does not determine motor output. More relatedly to our system, locusts display consistent individual-level preferences for foreleg selection in visually-guided gap crossing (51), but it was not established whether this is coupled to preferential eye use. Our findings differ in that both the sensory and motor components of the probing behaviour are lateralized and covary throughout continuous control, establishing a coupled sensory-motor axis rather than independent lateralized biases.

In hawkmoths, this lateralized sensory-motor axis was present from the first use of the proboscis and remained stable across days (Fig. 2A-C), indicating that the preferred control axis is established over the first contacts or is innate. At the same time, probing behaviour showed an experience-dependent refinement (Fig. 2D-F). This is consistent with previous findings in the tobacco hornworm hawkmoth that proboscis kinematics adapt with repeated interactions (60) and analogous to stick-handling crows, which improve their exploration trajectories with increasing experience (61). This suggests that while the directionality of the control axis may be constrained in lateralized moths, probing precision can be optimized through experience-dependent fine-tuning.

### Stability of the control axis under perturbation

Monocular occlusion of the frontal visual field revealed that neither the sensory nor the motor component of hawkmoth proboscis guidance showed significant flexibility to shift sides upon perturbation of the preferred side (Fig. 5A-B). Yet, the viewing angles of moths post-occlusion did not show a significant shift towards the lateral visual field of the occluded eye (Fig. 5J-K). We attribute this to the fact that the head keypoints of moths were closer to the stripe overall (Tables S6; Fig. 5C), which would bias the calculated mean viewing angles towards 0° (see *Methods*), hence the directional but non-significant effect. However, the significant shift in joint angles for both groups (Fig. 5G-I) could only be achieved by a combination of shifting in lateral body position towards the opposite side of the stripe (Fig. 5C), as well as a rotation of the head such that the lateral visual field becomes closer to it (Fig. 5D-F). Thus, considering the combination of body and postural adjustments, our data support that moths maintained their preferred control axis using the unobstructed lateral visual field.

This resulted in strengthened coupling between viewing angle and proboscis angle (Fig. 5N), indicating that the visuo-motor axis was maintained even when sensory input was degraded (Fig. 5D- E, trend in F). Thus, compensation occurred at the level of posture and flight control rather than by remapping the appendage to the opposite visual field. A similar trend was observed for moths with their non-preferred eye painted (Supplementary Fig. S5E-F, H-O). The persistence of the axis also extended to visual feature targeting, as moths continued to contact the preferred pattern edges after occlusion, despite the change in body position (Supplementary Fig. S5D). This indicates that the entire behavioural strategy guiding flower probing – from pattern feature targeting to proboscis guidance – is lateralized. Together, these results suggest that neither the sensory nor the motor component of the behaviour is independently dominant, and instead, the control axis itself appears to be the unit of lateralization. Such organisation may be advantageous for behaviours requiring rapid, continuous feedback, as in proboscis probing while hovering (43), where preserving a stable coordinate system could reduce the need for repeated re-calibration (62,63).

The persistence we observed in the visuo-motor axis contrasts with other insect systems, in which motor lateralization can be modified. While it has not been explicitly tested whether lateralized locusts can change motor output upon sensory perturbation, they can flexibly use either eye and either forelimb for reaching tasks following monocular occlusion (40). Indeed, when locusts (40) and proscopids (41) are subjected to monocular occlusion, they switch control strategies to the foreleg visible to the unobstructed eye. However, in non-occluded conditions for both species, both forelegs were used with equal probability – contrasting with the hawkmoths, which displayed preferred eye- proboscis coupling even before occlusion.

Moreover, locusts have lateralized motor circuits underlying their foreleg control (64), but it has not been shown how this reflects in motor behaviour. Additionally, motor preferences such as jump direction are pliant to experience (59), reinforcing the notion that they lack a rigid control axis guiding movement (or appendages more specifically). In hawkmoths, by contrast, the preservation of the control axis suggests that lateralization is embedded in the circuits linking visual input to proboscis motor output, rather than emerging solely from asymmetric eye use or motor biases. This distinction may reflect organization necessitated by different control tasks: discretized tasks like turning or leg extension for walking are typically performed in succession and often need to switch control repeatedly across the body midline. Thus, to enable flexible coordination for both locomotion and appendage placement, greater flexibility may be required in how (and which) legs and eyes are used. Conversely, continuous motor tasks like reaching typically guide a single appendage throughout its duration, and as such would benefit from a unified control axis rather than continuously remapping across both sides of the midline. Then, if only one appendage is required for continuous action, as in the hawkmoth proboscis, the rest of the body could adjust and compensate when needed to support it.

During similar continuous motor tasks, a small fraction of humans, chicks, and pigeons have been shown to reinforce the initial control axis (10,34,65). Hawkmoths also overwhelmingly retained this lateralized axis following occlusion, as well as the probing performance across all measured parameters (as in Fig. 3). This suggests that the perturbation strategy they use is beneficial to enable continued probing. Taken together, the general control required for continuous motor tasks could promote such reinforcement. However, most individuals in those vertebrate models tested show a complete switch of the control axis to the other side of the body, sometimes accompanied by performance costs. This flexibility suggests that continuous task structure alone might not be the driver of reinforcement per se, but perhaps the combination of task structure and invertebrate neural organization leads to the results we found. Moreover, how the control of paired versus unpaired appendages factor into this plasticity contributes to a series of intriguing future directions for comparative investigation between control strategies.

The reason why hawkmoths in our experiments did not switch control to the non-occluded side could also be that we blocked only the acute frontoventral region of the eye, preserving other visual functions in the lateral eye regions, such as wide-field motion-based flight control. Thus, we cannot rule out that an occlusion of the entire eye might also have revealed plasticity to shift control. Future whole-eye occlusion experiments could address this question, with the challenge of disentangling the impact of visual proboscis control and general flight control on task performance.

### Implications for neural implementation

The stability of the visuo-motor axis under partial monocular occlusion suggests that lateralized control is likely implemented at the level of neural circuitry, rather than arising from transient sensory dominance. Whether such lateralization and stability in hawkmoths arises from developmental sensory biases or intrinsic motor asymmetries remains unresolved, but provides intriguing directions for future investigation.

The high correlation between proboscis angle and pattern viewing angle we observed (Fig. 4G) suggests precise retinotopic alignment, as this analysis is based on the very strict assumption that the proboscis and pattern are viewed at the same visual angle off the midline for sensory-motor control. However, assuming wider visual fields of the underlying control neurons, as are typically found in other visually guided tasks (66,67), retaining pattern and proboscis within the same coarse visual field might be sufficient for control. Our data are consistent with this, as hawkmoths indeed retained their proboscis within congruent visual hemispheres 64% of the time in our experiments (Fig. 4G). Thus, the visuo-motor axes could operate through broader sensory-motor associations that preserve relative geometry between appendage and target. The range of coupling strengths across individual hawkmoths (Fig. 4H) suggests that there is likely not a single optimal neural mapping to achieve this, but that it depends on an individual’s sensory-motor system.

Subtle differences in musculature or pre-motor circuitry of the proboscis (44) could bias early motor output, which may then consolidate biased sensory-motor pathways. Another possibility is that visual pathways projecting to proboscis motor centres are asymmetrically weighted, forming lateralized sensory-motor loops that are similarly strengthened through Hebbian plasticity (68,69). Both possibilities would suggest that hawkmoths optimise their behavioural strategy within the constraints imposed by their sensory-motor system, giving rise to the lateralized visuo-motor axis. This optimisation would explain why we observed such consistent individual-level differences between hawkmoths (Fig. 1I-J), even within animals sharing the direction of lateralization. Consequently, unbiased animals may possess less restrictive constraints compared to their lateralized counterparts, allowing them both more flexibility in strategy, as well as possibly affording greater control during flower interactions.

However, distinguishing among these possibilities will require anatomical and physiological mapping of the circuits underlying proboscis guidance, combined with developmental and sensory perturbations. Indeed, neural investigations would confirm whether the individual-level variability we observed here is due to neural circuit asymmetries, which are linked to behavioural individuality in other insects (70,71). Such work would be instrumental to distinguishing the origins of sensory-motor plasticity.

### Functional implications of lateralized visuo-motor control

Lateralization has often been proposed to enhance behavioural efficiency by allowing neural circuits to specialise and reduce computational demands during complex tasks (72–78). In our paradigm, we did not find any evidence supporting this: stronger lateralization was associated with reduced probing precision and increased trajectory variability (Fig. 3E; Fig. 2F), suggesting that strong coupling comes at a cost of reduced degrees of freedom in maneuverability, due to the rigid lateralized axis. As there is strong evidence that pollinators, including hawkmoths, use patterns as nectar guides (43,45,79–82), we could expect that targeting patterns with maximum precision should lead to a fast nectar discovery. The longer probing durations and trajectory lengths (Fig. 3G-H) may thus indicate compensatory strategies that offset reduced precision through increased exploratory effort. As a result, strongly lateralized hawkmoths may experience increased foraging cost in time and energy.

Ecologically, such trade-offs likely influence how animals interact with complex environments. A consistent lateralized strategy could reduce decision time when approaching symmetric targets (83), facilitating rapid interaction with flowers by breaking symmetry, potentially reducing energetic costs of flight (43,45,55,79–82). However, we did not find this trend in our data. Lateralized hawkmoths had higher flight speed during the approach phase, but the overall duration of the approach did not differ from unbiased individuals – presumably an effect of compensation (Supplementary Fig. S3G-H). Slower speed during this phase in unbiased moths could indicate a more stable, controlled approach from proboscis unfurling to the initial flower contact, later reflected in the increased precision (Fig. 3E).

Flower probing proves a challenging control problem for hawkmoths – maintaining a stable hovering position in front of flowers, guiding the proboscis, and monitoring the horizon for stability and predators. The demands of flower probing are compounded when facing unfavourable conditions such as monocular occlusion. The strengthening of visuo-motor coupling under monocular occlusion (Fig. 5L-N) could therefore suggest that increased task difficulty promotes stronger reliance on specialised control pathways (‘Task Complexity Theory’, (84)). Whether lateralized control strategies are favourable for adapting to challenging sensory conditions, and how this impacts foraging success and energetic efficiency in natural environments, remain open questions that can now be addressed experimentally in the hawkmoth proboscis model. For now, the functional consequences of lateralization appear context dependent in hawkmoths, similar to the performance consequences of lateralization in other taxa (85). Our work argues for lateralization as a form of behavioural optimisation under specific sensory and motor constraints, rather than a universally beneficial strategy.

## Conclusion

Together, these findings demonstrate that continuous sensory–motor coordination can be organized around a conserved lateralized axis with flexible flight compensation. Our work thus provides a general framework for understanding how lateralization shapes control strategies enacted by compact nervous systems. It can be directly compared to the lateralized axis found in human, elephant, and avian appendage guidance, highlighting convergent principles, as well as species- and context-specific differences in plasticity and performance consequences of lateralization.

## Material and Methods

### Aninals

*Macroglossum stellatarum* (Sphingidae) males and females were taken from lab colonies at the University of Konstanz, Germany. Adult animals were kept in flight cages (60cm x 60cm x 60cm), with a 14:10h light:dark cycle at 23°C. Adults could feed *ad libitum* via gravity-assisted feeders filled with 20% sucrose-water solution (86). The ‘feeding-experienced’ adults used in these experiments were between 1 to 6 weeks of age. Prior to experimental trials, randomly selected adults were separated into marked plastic vials and kept in the dark for approximately 24 hours to ensure no further feeding occurred. Naïve animals were the result of spontaneous matings within the lab colony, where multiple female *M. stellatarum* laid egg batches on the host plant *Gallium sp.* Upon hatching, caterpillars were reared on the host plant until pupation (approx. 3 weeks). Following eclosion (pupal phase lasting approx. 3 weeks), fresh adult moths were kept in a hatching cage without food for approximately 24 hours to allow wings to fully unfold and to take flight for the first time, before being transferred to plastic vials and marked as ‘naïve’. These naive moths were then tested the following day.

Additional data were used from an existing dataset of a previous study ((43); data repo: https://doi.org/10.6084/m9.figshare.22639981.v1).

### Experimental Setup

The experimental setup and paradigm of this study are largely based off that shown in (43), and will be described here in brief.

Experiments were conducted in a cubic metal flight cage spanning 60cm x 60cm x 60cm at 25°C and 30-45% humidity. The cage was illuminated from above by four 60cm long daylight-like fluorescent tubes (Osram L 18 W/965 Biolux Tageslicht G13). All lights were connected to an electrical ballast (GloMat 2 x 40 W, Hagen), which increased the flicker frequency of the fluorescent tubes above 25kHz, outside of the resolvable range for hawkmoths. Two infrared light arrays were also placed above the cage to enhance visual contrast in video recordings, at wavelengths which hummingbird hawkmoths do not perceive (>650nm; 87). The walls of the cage consisted of thin medical gauze, such that air and light could penetrate, but the animals could not leave. Resting above the cage, but beneath the lights, were interwoven nets of red twine, which provided visual contrast for optic flow and distance estimation in the dorsal view of the moths, and thus prevented the moths from flying into the cage ceiling. A 25-cm high pedestal coated with white felt was placed in the center of the flight cage, on which the artificial flowers were mounted. A camera (acA1300-200uc, Basler) was mounted directly above the ceiling of the flight cage, focusing on the artificial flower in the center of the cage. The IR-stop filter in the camera was removed prior to mounting. It was controlled using the Pylon Viewer Software (Basler) at 200 frames per second (played back at 30fps), with a field of view of 496 x 500 pixels (ca. 12 x 12 cm in the focus plane), recording in Mono 8 format (grayscale).

### Stimuli and Visual Patterns

All artificial flowers were constructed from round paper cut-outs of 38 mm diameter. All stimuli were printed on unbleached paper (“Classic White”, Steinbeis) using a laser color printer (imageRUNNER ADVANCE DX C3926i, Canon). All stimuli used in this study had a blue background, with a yellow stripe pattern in the center. The stripe had a width of 2 mm, and was 38 mm long, extending over the entire diameter of the flower. The yellow colours were set as C=0%, M=0%, Y=100%, and K=0% and the blue as C=91%, M=26%, Y=0%, and K=0%. They were laminated with a matte foil (S-PP525-22 matte, Peach, PRT GmbH) after printing and trimming and cut out to shape again.

### Experimental Procedure

In this study we performed three sets of experiments, in which the stripe pattern was shown to moths of two groups of feeding experience: naïve and experienced. Feeding experienced moths had fed from gravity feeders (86), but not been in contact with the artificial flower stimuli used in our experiments. In the first experiment, each experienced moth was allowed to freely fly in the experimental setup containing the pattern stimulus, for a 10-minute period. The artificial ‘flower’ was presented without sugar water or odour cues during this period. Moths could visit the artificial flower an unlimited number of times within this period. All approaches and probing phases were recorded with a manual trigger. If a moth probed the flower with its proboscis, it was rewarded at the end of the 10-minute period ad libitum with 20% sugar solution on a feeding disc. The disc was identical to the pattern used during the 10-minute period, and temporarily replaced the initial stimulus on the pedestal. This aimed to minimize the possibility of odour or gustatory cues from sticking to the experimental stimulus. If a moth did not probe the pattern within the 10-minute period, it was not fed at the end of the session, and packed in the holding container until the experiment was repeated the following day. If an animal did not approach within two consecutive days, it was excluded from the experiment and returned to the communal feeding cages. In between testing days, all animals were held in individual holding tubes, and placed in a dark container until the next session began. The flower pattern was rotated in their horizontal orientation by 90° between subsequent animals and days, to not be presented in the same orientation relative to the surrounding cage. Experienced animals were tested for 3 to 5 consecutive days before either being returned to a tested animal cage, or kept for the second experiment.

The second experiment was identical to the first, except naïve animals were tested for up to five consecutive days, depending on motivation. After the final day of testing, animals were then returned to a communal flight cage and could feed *ad libitum* from feeders.

The third experiment began with analyzing the proboscis lateralization of experienced animals across testing days. Then, animals were assigned to a distinct subgroup: left-lateralized moths showing a significant leftwards proboscis direction in their overall distribution of bout positions; right-lateralized moths showing a significant rightwards proboscis direction in their distributions; and unbiased moths showing an alternation in proboscis placement preference across days (e.g. *LRLRL*), or placement positions being non-significantly different from zero. The fronto-ventral area of the hawkmoths’ corresponding eye (e.g. left eye for a left-lateralized moth) was painted to create two treatment groups: left or right eye painted (experimental group split based on left/right lateralization groups). See *eye painting* methods below (Supplementary Fig. S5*A*). The animals were then allowed a 2-hour period to acclimate to the paint, by being placed in a plain flight cage individually. Following the acclimation period, moths were returned to holding tubes and kept in the dark until the following day. Painted moths were tested for three more consecutive days in the experimental setup, or until motivation ceased, under the same regime as the non-painted animals. The animals were collected after this three-day period, and their eyes were photographed from several orientations using a Flexacam C1 (Leica) camera mounted on a M80 stereomicroscope with 10 x oculars and a 1 x objective (Leica).

### Eye Painting

For the third experiment, individual moths were briefly anesthetized with a 3-4s exposure to concentrated CO_2_ in their holding tube. During this anesthetized period (approx. 45 – 80sec), a black water-based lacquer was painted onto the fronto-ventral region of the eye using a small brush, corresponding to the proboscis lateralization direction (Supplementary Fig. S5*A*). The painted region thus extended approximately from the equator of the frontmost portion of the eye, ventrally by about 0.7mm. The lateral boundary of the painted zone followed an arc which peaked approximately 0.5mm from the innermost edge of the eye.

### CO_2_ Control Testing

It was previously shown in (43) that brief anesthesia and painting single eyes did not disrupt the ability of moths to freely fly and interact with the pattern stimulus. We further validated this by briefly anesthetizing a group of non-experimental moths with the same protocol, and observing their behaviour in a communal feeding cage. Within 20 minutes, all moths were able to freely fly, interact with the feeders, and perform reproductive behaviours such as mating and laying viable eggs. Additionally, we performed the same protocol on a small sample of naïve moths following the second experiment, anesthetizing them and testing their probing behaviour the following day, which confirmed that they showed similar behaviours as before the treatment.

### Video Tracking

We used DeepLabCut (ver. 3.0.2, (88)) to automatically identify the proboscis tip, head tip/proboscis base, head centre, antennal bases, and thorax position of the hawkmoths (see Fig. 1*C* for skeleton). A PyTorch model was trained using 1792 manually labelled frames of hawkmoths in various orientations, using the HRNet_w32 architecture (89) and associated ‘cutout’ method (90). 95 frames were randomly withheld from the training dataset to be used for testing (∼5% of frames). The model was trained for 200 epochs, resulting in a training RMSE of 0.82 pixels (without p cutoff), train mAP of 99, train mAR of 99.74, test RMSE of 1.02 pixels, test mAP of 98.21 and test mAR of 99.05. Tracked keypoints were passed through DeepLabCut’s in-built filtering process to minimize jitter. Despite the model’s performance on the test dataset, there were still frequent small gaps of data missing, particularly for the proboscis tip. All keypoints were then passed through a modified Akima interpolation filter in custom-written Matlab code to fill in gaps of <= 5 consecutive frames.

All videos were manually corrected for accuracy of the filtered and interpolated tracking points, particularly of the proboscis tip, which was still occasionally subject to inaccurate tracking after filtering, using the DLTdv8a software (91) in Matlab 2024a (The Mathworks). Only proboscis coordinates that were tracked when the proboscis was in contact with the artificial flower in the current frame were retained. Contact with the flower surface was noticeable in the small bend of the proboscis on its very tip. Using the same software, the flower position was marked by four points on the flowers’ circumference parallel to the cardinal axes, and the pattern position was marked on all pattern edges in each video. Data were then further processed using custom-written Matlab scripts.

## Data Analysis

For each animal (total number given as *n* in the figure legends), tracking data were generated for the head, proboscis base, thorax, and proboscis tip positions for each experimental session (across all three experiments). Within a session, the hawkmoths typically approached the flower multiple times, with varying lengths of probing times (and therefore number of contacts). Each of these individual approaches was registered as a trial or ‘bout’ (number given as *N* in figures and figure legends, unless otherwise indicated), which consisted of the approach flight (250 ms before the first proboscis contact), the probing phase (continuous proboscis contacts with the flower without interruption for more than 20 ms), and the departure phase (250 ms after last contact). Only animals with at least 500 proboscis contacts across all trials (corresponding to a total probing time of 2.5s per condition) were included in the analysis. Additionally, bouts with less than 30 contacts were excluded from analysis.

Before further processing, all tracking points (of the hawkmoths and the flower) were rotated so that the long axis of the pattern was aligned with the y-axis of the Cartesian coordinate system used throughout (Fig. 1*C*). Thus, the data were transferred from a cage-centric to a pattern-centric coordinate system. Positional data were converted to world units using a pixel-millimeter scale factor.

## Standardised bouts

Animals in these experiments generated their own bouts, with variable duration and frequency. For analyses concerned with comparing proboscis distributions and proboscis variability between individual animals (Fig. 1, Fig. 2), we standardised the number of contacts within each bout. To do so, we calculated the median bout duration across all animals (127 contacts). Then for each animal, we found all contacts and segmented them into standardised bouts by the median bout length. The resulting number of bouts varied across animals, but contained equal numbers of contacts. Standardised bouts closely resembled those of the animal-generated bouts (Fig. S1*B*; r=0.95, p<0.001; Fig. S1*C*; r=0.94, p<0.001).

## Proboscis position

To determine relative proboscis position, an axis between the head centre and the proboscis base (tip of the head; ‘PB’) served as a moth-centric reference frame (Fig. 1*C*). Then for each proboscis contact, the proboscis base position was subtracted from the proboscis position to establish relative x/y coordinates and stored in a matrix. The x-position data were also summed in bins spanning - 15 to 15 mm relative to the moth’s head, to be used for density distributions. Relative angle of the proboscis to the head axis was calculated by first finding the heading direction of the head-PB vector. Then, the head axis and PB-proboscis vectors were normalized and used to calculate the dot product. Relative angle was found by taking the arc cosine of the product, with the determinant of the two vectors representing the sign of rotation, summarized in **Equation 1**.

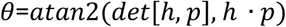

Heatmaps of the relative proboscis positions were generated by computing the normalized mean density of each animal, then applying a Gaussian filter (sigma=2.5) to the normalized data. Contours within heatmaps were generated by binning positions with 0.1-unit steps from 0 to 1 normalized density. Density distributions of the proboscis lateral x-positions were computed through normalizing the summed x-positions by the sum under the curve. The data were then scaled by the maximum value of the normalized distribution. The maximum of the distribution was used to calculate the lateral x-position with the largest density of proboscis position (per-bout and overall).

## Proboscis contacts on the flower

To depict the distribution of proboscis contacts on the artificial flower pattern, the positions of the proboscis of each animal in all trials were collected in a matrix, which represented the spatial layout of the flower. To compare animals with different numbers of proboscis contacts on the flower, contacts were summed for each animal, and then divided by the total number of contacts per animal. These normalized contact distributions were then averaged across animals, to generate the proboscis contact heatmaps.

## Lateralization direction and lateralization index

We calculated two metrics for determining the strength and direction of proboscis lateralization using standardised bouts. The first uses world units (millimeters), consisting of the maximum of the proboscis position distribution relative to the head, for each of the animals’ bouts, referred to as lateralization ‘amplitude’. This resulted in a distribution of per-bout maximum proboscis positions per animal. These distributions were than statistically compared to zero to determine lateralization direction (negative=left, positive=right, zero=unbiased), with distribution medians representing the amplitude for each animal.

We also calculated a modified lateralization index (LI) per-bout and overall, matching the sign of world units (left=negative, right=positive), as depicted in **Equation 2**.

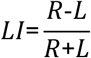

R represents the number of contacts to the right of the animal’s head axis (positive), and L represents the number to the left (negative). The output is a ratio indicating lateralization direction (left=negative, right=positive), alongside relative lateralization strength (-1/1 indicating strong lateralization; 0 indicating no lateralization). Similar to the previous metric, these distributions were also compared to zero, per-animal, to determine significance of the lateralization indices observed. As lateralization index and median bout position were closely matched (Supplementary Fig. S1*C*), we used median bout position as a measure of lateralization amplitude and direction for subsequent analyses.

## Bootstrapped distributions and model fits

Bootstrapped distributions of proboscis lateralization direction, amplitude, and interquartile range (IQR) were performed. First, we calculated the median number of bouts within days, across animals (median=11). Then, we resampled with replacement the bout positions for 10,000 iterations, by the median number of bouts per simulated animal. From each resample, we counted the number of animals within each lateralization category and calculated the average number for each group across all resamples. We also calculated the median maximum bout position and IQR of bouts per simulated animal within each resample, totalling 10,000 of each measurement per animal. From this, we computed the median bout position and IQR to compare directly with that of the observed data.

## Bout positions per day

To represent how animals’ median bout-wise positions changed across days (individually and group-level), we calculated the difference in each animals’ median across consecutive days using standardised bouts. We then calculated the mean difference in medians at a group level, across consecutive days. Estimation statistics were used to determine the significance of these changes and acquire a measure of group consistency (92,93). Only animals with at least 3 consecutive days of testing, containing a minimum of 5 bouts, were used in this analysis. Visualizations in Figure 2 depict the first 3 days, but subsequent variability analyses include all days of testing (all consecutive days in Supplementary Fig. S2*A*).

## Variability as interquartile range

Given the non-normal distribution of maximum bout positions within and between animals, we used interquartile range (IQR) as a non-parametric measure of dispersion (and thus variability). We used two measures of IQR to determine consistency and variability of proboscis lateralization. The first measure was within-day variability, representing how variable an animal’s proboscis positioning is within a day. To calculate this, we found the IQRs of bout positions for each of the days an animal was tested, resulting in 3 to 5 IQR values per animal. Then, we used the median IQR value as the proxy for within-day variability for each animal. The second measure was across-day variability, or how much a moth tended to shift its overall proboscis lateralization across days. Conversely to the within-day measurement, we calculated the median bout position for each of the days an animal was tested, providing 3 to 5 bout medians per animal. Then, we calculated the IQR of these bout medians as a measure of how variable an animals’ central tendency was across days.

## Relative head orientation

We used the axis between the head centre and proboscis base as the reference for head orientation. Using a polar coordinate system (-180°,+180°), where clockwise (CW) depicts negative angles and counterclockwise (CCW) indicates positive angles, we fixed the head centre as the axis origin (see Fig. S4*B*). We further rotated the coordinate system by 90° such that 0° points upwards. All postures were rotated such that if the animal was in the upper half of the frame, its head orientation was now positioned in the lower half with the same relative angle, such that facing upwards (parallel with the stripe) indicated 0°. We then mirrored any angles facing downwards (>90°, <-90°), constraining all orientations between (-90°, +90°]. The relative angles of all animals were first pooled to produce a group polar histogram distribution. Then, the circular means, 95% confidence intervals (CI), and mean significance of heading angles were extracted per-animal for statistical analyses.

## Relative joint angle orientation

Following calculation of the head angle, we calculated the relative orientation of moths to the center of the pattern ([x,y] = [0,0]). Using the relative head vector, we then set the proboscis base to be the origin and calculated the relative joint angle between this vector and the pattern center (see Fig. S4*E*). Using methods described for calculating relative proboscis angle, we calculate the angle as seen in **Equation 3**. Angles which were outside the interval [-90, 90]° were rare and omitted from the calculations. Similarly, this produced a distribution for the group, as well as per-animal means, 95% CIs, and significance.

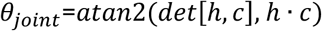

## Occupancy heatmaps

We constructed the density heatmaps of moth head positions by creating a 500x500 matrix for each animal, and summing the occupancy of the moth’s head in each cell across the entire video. Then, the occupancy matrix for each moth was normalized by the .999 quantile before calculating the mean occupancy of all the normalized matrices. The resulting group occupancy map was smoothed with a Gaussian filter (sigma=2.5) and normalized for color scaling. This created a normalized relative occupancy map (0 to 1) for moth head positions surrounding the pattern stimulus.

Difference occupancy maps represent the difference in positional density between two groups, or two conditions. First, each group’s density was computed similar to the previously described map and smoothed with Gaussian filters. Then, the resulting difference map was computed by subtracting the first map from the second. This was again normalized by the .999 quantile, such that negative values indicate a greater relative occupancy by the second group, and positive values indicate greater occupancy by the first group.

For comparing occupancy differences before/after painting (Fig. 5C), we only analyzed data where moths were oriented parallel within [-45°,45°] of the stripe, to assess how their lateral positions changed in response to occlusion.

## Average body part positions relative to pattern

To compute how the animals’ body was positioned relative to the pattern during probing bouts, we computed the average positions of three body parts per animal-generated bout. We first normalized the positions of the proboscis tip, PB, and head relative to the midpoint of the stripe, such that the center of the stripe is zero, leftwards is negative, and rightwards is positive. The parallel edges of the stripe were thus positioned at -1mm and 1mm. Then for each bout, we calculated the average x- position of each body part relative to the stripe, and transformed it into world units using a pixel scale factor. Thus, we produced relative mean body part positions for each of the animals, as well as for the group overall. For proboscis contact density distributions, the proboscis contact positions were summed within each bout in bins of 0.5mm spanning the width of the stimulus. Then for each animal, we computed the mean binned x-position distribution, and normalized by the maximum density to acquire a per-animal contact distribution. To acquire a group-level contact distribution, the distributions of all animals were pooled and averaged to be normalized in a similar manner to the per- animal distributions.

For comparing body positions before/after painting (Fig. 5C), we only analyzed data where moths were oriented parallel within [-45°,45°] of the stripe, to assess how their lateral positions changed in response to occlusion.

## 2D visual field vectors

We approximated the moths’ visual field from the top-view (2-dimensions; x/y) by generating rays which extended out from the midpoint of the head, and thus passed through the location of the eyes. First, we created rays of 715 pixel length, approximately the diagonal length of a frame. Then, we separated rays into 1° bins from [-110°, 110°], simulating the finest facet acceptance angle (94), while assuming no binocular overlap between the eyes. To organize these rays into left/right visual fields, the rays were rotated using a polar coordinate system such that the left visual field consisted of rays pointing from [110°, 1°] and the right consisting of rays facing [-1°, -110°], relative to the heading direction. **Equation 4** depicts the rotation calculation for each ray, per frame.

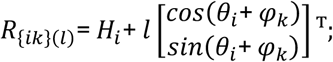

To determine relative distance of a ray to the pattern, we used the stripe’s corner coordinates to generate a rectangular polygon. Then, per-frame, we calculated the relative length of each ray in Euclidean distance. If in contact, the length of a ray was truncated by the edge of the polygon, effectively shortening the relative ray length compared to the default length. Thus, we could calculate the following: shortest ray length and the angle of the shortest ray. Frames where a moth’s head midpoint was located inside the polygon were not included in analyses, as all rays would effectively be the shortest length.

## Mean viewing angle

In addition to calculating the shortest length, we also calculated a mean ray angle, which approximated the weighted mean viewing angle of the moth to the pattern. To do this, we first calculated the inverse distance to the polygon by subtracting each ray’s length from the default length, effectively weighting the rays by proximity to the stripe. Then, we normalized all ray lengths by the default ray length, scaling the vectors of each ray from 0 to 1. Thus, any rays not in contact were equal to zero, whereas rays with shorter distances to the polygon approached a value of 1. This simulates closer objects having a larger relative visual angle, and therefore potentially providing more information or having more influence in the visual system. We then converted from polar to Cartesian coordinates by multiplying the normalized ray lengths by their respective sine/cosine components, as depicted in **Equation 5**.

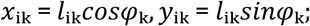

This allowed us to calculate a mean vector of all intersecting rays, by finding the mean x/y components of all rays, and convert the components back into a resultant mean vector angle and length through **Equation 6** and **Equation 7**.

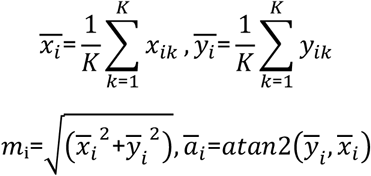

Thus, per-frame, we generated a weighted mean pattern viewing angle and length, as well as the relative distances of each ray to the stripe.

## Proximity-weighted intensity of visual scene

To simulate the approximate visual scene of the animals, we transformed the per-frame proximity-weighted lengths of each ray into a relative intensity. Therefore, rays with a value closer to 1 represented a higher relative intensity, and a closer proximity to the stripe. The output of this calculation is a distribution of relative intensities per-frame, per-animal, effectively generating 1- dimensional visual scenes. We used these distributions to calculate the mean intensity across the visual field over all frames, resulting in a mean intensity distribution per animal. Using the mean intensity distributions of all animals, we calculated a smooth mean intensity distribution with a moving average (local k-point=5). We further extracted the peak of each animal’s mean intensity distribution as an estimation of which visual angle was used most often to view the pattern, or had greatest relevance throughout the pattern interactions.

## Bout position changes after painting

To determine whether the relative proboscis positioning changed after animals had their eye painted, we calculated the difference in median bout position before/after this treatment. First, we pooled all bout positions within each animal and calculated the median as a measure for ‘before’ painting treatment. Then, we performed the same calculation for the bouts recorded ‘after’ painting treatment. The difference in median was calculated per animal, as well as the average median bout position of each group in both treatment conditions.

## Proboscis-pattern edge differences

To calculate the difference between proboscis placement and an edge of the stripe pattern, we first found the average position of the proboscis relative to the stripe (within each bout). Then, we computed the difference between the absolute value of this position, and 1mm, which corresponded to the absolute position of either edge. Negative values indicate average proboscis positions inside the stripe (between [-1,1]mm).

## Full-width half-max of contact distributions

As a measure of precision, we computed the full-width half-max (FWHM) for each bout’s proboscis contact position distribution, across animals. To do this, we first found the x-position of the distribution maximum. Then, we found the x-positions of the distribution which corresponded to half of the maximum, for both sides of the maximum. The distance between these two midpoints was calculated as the FWHM. Bouts containing multiple peaks of the same value rarely occurred, and the maximum position was computed as the average of the peaks’ positions.

## Bout interaction parameters

Within each bout, we computed parameters relevant to the kinematics of the probing trajectory. Bout duration was computed as the time elapsed across a probing bout, converted from frames to seconds. Track length was measured as the total distance covered by a bout’s probing track. Track tortuosity was calculated as the total length of the track, divided by the Euclidean distance between the track start and end point, with lower values indicating straighter paths. Probing speed was computed as the median Euclidean distance between the consecutive points of a track, divided by the framerate of the camera. Time taken from when a moth entered the video frame to first proboscis contact was used to calculate approach duration (sec) and approach speed (mm/sec). Duration was calculated as time taken between frame entrance to first contact, while speed was calculated from the median Euclidean distance between consecutive points of the head during approach, divided by the framerate of the camera.

## Statistical Analysis

Statistical analyses were performed using custom scripts written in R (v. 4.4.3) using RStudio (v. 2024.12.1), MATLAB R2024a (The Mathworks), and through the Python-based estimation statistics toolbox ‘DABEST’ (93) in Python v. 3.11.0. Relevant statistical tests were chosen based on properties of the data distributions and normality assumptions. Normality assumptions of distributions were tested using the Shapiro-Wilk test. For unpaired, independent tests between distributions, the non-parametric Wilcoxon rank-sum test was used. For paired distribution tests, or tests between distributions and zero, the non-parametric Wilcoxon signed-rank test was used. Paired and unpaired tests were treated as distinct test sets and corrected separately for repeated testing biases using Hommel’s p-value correction method (95). For comparisons between >2 non-normally distributed groups, the Kruskal-Wallis test was used with Hommel’s correction. Comparisons between the shape of distributions were performed using the Kolmogorov-Smirnov test.

Circular data were analysed using the MATLAB-based ‘Circular Statistics’ toolbox (96). Departures from uniformity were assessed with Rayleigh’s test. All other statistical analyses on circular data were performed using functions from this toolbox, including: circular mean, circular median, circular mean confidence intervals, V-test for nonuniformity in a specified direction.

Estimation statistics were performed using in-built functions of the DABEST toolbox. Effect sizes were calculated from two-tailed permutation tests on 5000 reshuffles of the data, with 95% confidence intervals calculated from the bias-corrected and accelerated distribution. Effect sizes were reported as either mean differences, or as Hedges’ g for paired comparisons below sample sizes of 20 individuals. Effect sizes were considered significant based on a lack of overlap between the intervals and zero, further supported by p-values calculated for the effect.

When applicable, bespoke statistical models were used to determine estimates and significance of relevant parameters in R:

Linear regressions were performed with Matlab’s *corrcoef* function, the R function *cor.test*, and R ggplot2 function *scatter.smooth(method=’lm’*) (97). All correlational analyses performed with linear models report the Pearson correlation coefficient.

To confirm that within-day variability differed between naïve and experienced moths, we used a generalized linear mixed model fitted with the glmmTMB function (98) on the within-day IQR data. We initially included all predictor variables we found relevant, including experience group, day tested, the absolute per-day median proboscis position, an interaction between experience group and day tested, random slopes per animal, and random intercepts per animal. We sequentially dropped one factor at a time, comparing the reduced model to the full model, using the log likelihood ratio and relative AIC values to determine the better fit. From this, we found the best fit was provided by a formula containing experience as a fixed effect, absolute per-day median proboscis position as a fixed effect, day as a fixed effect, and animal ID as a random intercept. The best fit also included absolute proboscis position as a random slope, as different animals may vary in their positions over time, so we centered the absolute positions for both the fixed and random effect:

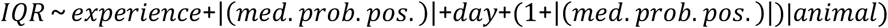

To account for the right-skewed distribution of data, we used the lognormal family with a log link. We simulated residuals using the DHARMa package ((99); v. 0.4.7) to validate our model’s goodness of fit. Log estimates and profile confidence intervals were backtransformed from log to original units using an exponential function.

A generalized linear model (glm, R ‘stats’ package v. 3.6.2) was fitted to the across-day IQR data. As there was only one across-day IQR measure per animal, we kept the same model structure but dropped day as a fixed effect, and animal ID as a random intercept, while using the overall absolute median proboscis position:

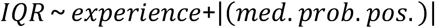

The Gamma family with a log link was used to fit the data, due to a slightly different skew in response distribution. Goodness of fit was assessed by simulating residuals with the DHARMa package. Log estimates and confidence intervals were backtransformed using an exponential function.

Similar models were fit to determine how lateralization strength (LI) affected variability. Within-day variability was fit using the below model, after centering the absolute lateralization indices. Including day as a factor did not improve the fit, so it was dropped. The response distribution did not change, so we kept the lognormal family with the log link.

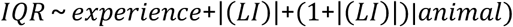

Across-day variability was fit using the below model, again swapping out proboscis amplitude with lateralization strength. The rest of the model remained the same.

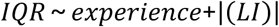

To determine whether the bout-wise average position of the three body parts differed amongst lateralization subgroups, as well as before/after monocular occlusion, we used a robust linear mixed model to fit our data (*rlmer*, (100)). This robust model automatically adjusts weighting of outlier residuals and datapoints to reduce the influence of extreme values, particularly for hierarchical and/or nested data. The structure of our data included three nested measurements (body parts) within each bout per animal, representing a nested dependent sequence of datapoints. The model included a fixed interaction term between body part and lateralization (e.g. left/right/unbiased), as we sought to determine whether each group positioned themselves differently. We used 0 as a fixed effect, removing the estimate for an overall intercept, instead estimating the means of each combination of body part and subgroup. Robust models could not converge when including body part as random slopes, due to the number of random effects needed to fit the data. However, we included bouts nested within animals as a random intercept, allowing the intercepts to vary within each bout per animal. The resulting formula was used:

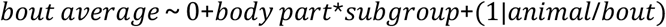

For analysis of the experienced animals before/after occlusion, we swapped out ‘subgroup for ‘eye condition’, comparing the difference between left-unpainted (LU), left-painted (LP), right-unpainted (RU), and right-painted (RP):

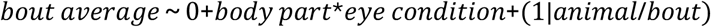

Due to downweighting and difference in model structure, typical methods for acquiring confidence intervals and assessing goodness-of-fit were not applicable (100). Instead, we acquired significance of estimates and contrasts by cluster bootstrapping the model fit for 5,000 iterations, calculating the 95% confidence interval of each estimate (and contrast) based on these resampled fits. If an animal was included in the resampled pool, its ‘cluster’ of data (i.e. all bouts and measurements within bouts) were included in the model fit. This allowed us to generate confidence intervals for estimates in the original model, and compute contrasts between the factor levels.

Detailed output of statistical models are available in Tables S1-S6. Details for specific statistical tests performed are described in figure legends and in the relevant Results sections. Full statistical test parameters are listed in Supplementary Tables S7-S12. Summary quantities (e.g. mean, median, etc.) used for statistical analyses are also indicated in relevant figures and Results sections. Significance levels are reported using the following convention: NS (not significant; p>0.05), * (p<0.05), ** (p<0.01), *** (p<0.001).

## Data, Materials, and Software Availability

Raw tracking data, example videos, and eye images (before/after painting) will be made publicly available upon publication. Detailed results of additional statistical tests used for analyses are reported in Supplementary Tables. All custom-written code used for analysis and visualization of data is available in the GitHub repository: https://github.com/stoeckl-lab/ProboscisLateralisation.

## Acknowledgenents

We acknowledge funding to A.S. by the Bavarian Academy of Sciences and Humanities, the Zukunftskolleg Konstanz, the Hector Fellow Academy, and the Emmy Noether Programme of the DFG (STO 1255/4-1). L.W. thanks the Hector Fellow Academy for support. This study was partially supported by the International Max Planck Research School for Quantitative Behaviour, Ecology and Evolution (IMPRS-QBEE). We are very thankful to James Foster for useful input on statistical models. We are grateful to Niels Poulsen for detailed input regarding optimizing DeepLabCut model parameters.

## Author Contributions

**Conceptualization:** L.W., S.K., A.S.; **Methodology:** L.W., S.K., A.S.; **Validation:** L.W., A.S.; **Fornal Analysis:** L.W., A.S.; **Investigation:** L.W., S.K.; **Resources:** A.S.; **Data Curation:** L.W., S.K.; **Writing – Original Draft:** L.W.; **Writing – Review G Editing:** L.W., A.S.; **Visualization:** L.W.; **Supervision:** A.S.; **Funding Acquisition:** A.S.

## Conpeting Interests

The authors declare no competing interests.

## Statistical Model Tables

**Table S1A.** – **Generalized linear nixed nodel** for the within-day IQR of experienced and naïve hawkmoths. Formula: iqr ∼ experience + abs. (median proboscis position) + day + (1 + abs. (median proboscis position) | animal), family=lognormal, link=log. Observation count = 236. Random-effects groups: animal=61. AIC = 706.4. Lognormal dispersion = 1.13. Log estimates are listed below.

|  | *Estimate* | *Std. Error* | *Z value* | *Pr (>\|z\|)* |
| --- | --- | --- | --- | --- |
| **Intercept** | **0.49528** | **0.08472** | **5.846** | **5.03e-09\*\*\*** |
| **Experience: N** | **0.31870** | **0.09766** | **3.263** | **0.0011\*\*** |
| **abs. (prob. pos.) centered** | **0.11418** | **0.04586** | **2.490** | **0.0128\*** |
| Day2 | 0.06810 | 0.08672 | 0.785 | 0.4323 |
| **Day3** | **0.15765** | **0.07773** | **2.028** | **0.0426\*** |
| Day4 | 0.06282 | 0.10429 | 0.602 | 0.5469 |
| Day5 | 0.01775 | 0.10922 | 0.162 | 0.8709 |

**Table S1B.** – **Backtransforned estinates and G5% CI for Model S1A** using the exponential function.

|  | *Estimate* | *2.5%* | *97.5%* |
| --- | --- | --- | --- |
| Intercept | 1.640950 | 1.3813016 | 1.928939 |
| Experience: N | 1.375335 | 1.1304177 | 1.669306 |
| abs. (prob. pos.) centered | 1.120959 | 1.0186247 | 1.223996 |
| Day2 | 1.070474 | 0.9017017 | 1.268999 |
| Day3 | 1.170751 | 1.0047014 | 1.365096 |
| Day4 | 1.064836 | 0.8621904 | 1.301613 |
| Day5 | 1.017906 | 0.8152724 | 1.735326 |

**Table S2A.** – **Generalized linear nodel** for the across-day IQR of experienced and naïve hawkmoths. Formula: iqr ∼ experience + abs. (median proboscis position), family=Gamma, link=log. AIC = 48.7. Nagelkerke’s R2 = 0.184. Gamma dispersion = 0.3791316. Log estimates are listed below.

|  | *Estimate* | *Std. Error* | *Z value* | *Pr (>\|z\|)* |
| --- | --- | --- | --- | --- |
| **Intercept** | **-0.7347** | **0.1377** | **-5.335** | **1.65e-06\*\*\*** |
| **Experience: N** | **0.4794** | **0.1739** | **2.757** | **0.00779\*\*** |
| abs. (prob. pos.) | 0.1853 | 0.0983 | 1.885 | 0.06445 |

**Table S2B.** – **Backtransforned estinates and G5% CI for Model S2A** using the exponential function.

|  | *Estimate* | *2.5%* | *97.5%* |
| --- | --- | --- | --- |
| Intercept | 0.479660 | 0.3685395 | 0.6317472 |
| Experience: N | 1.615095 | 1.1538149 | 2.2958874 |
| abs. (prob. pos.) | 1.203573 | 1.0040114 | 1.4589573 |

**Table S3A.** – **Generalized linear nixed nodel** for the within-day IQR of experienced and naïve hawkmoths. Formula: iqr ∼ experience + abs. (lateralization index) + day + (1 + abs. (lateralization index) | animal), family=lognormal, link=log. Observation count = 236. Random-effects groups: animal=61. AIC = 711.5. Lognormal dispersion = 1.21. Log estimates are listed below.

|  | *Estimate* | *Std. Error* | *Z value* | *Pr (>\|z\|)* |
| --- | --- | --- | --- | --- |
| **Intercept** | **0.56699** | **0.06427** | **8.822** | **< 2e-16\*\*\*** |
| **Experience: N** | **0.33641** | **0.09070** | **3.709** | **0.000208\*\*\*** |
| **abs.(LI)**<br>**centered** | **-0.37374** | **0.16387** | **-2.281** | **0.022568\*** |

**Table S3B.** – **Backtransforned estinates and G5% CI for Model S3A** using the exponential function.

|  | *Estimate* | *2.5%* | *97.5%* |
| --- | --- | --- | --- |
| Intercept | 1.762961 | 1.5542972 | 1.9996373 |
| Experience: N | 1.399917 | 1.1719134 | 1.6722793 |
| abs. (LI) centered | 0.688155 | 0.4991105 | 0.9488024 |

**Table S4A.** – **Generalized linear nodel** for the across-day IQR of experienced and naïve hawkmoths. Formula: iqr ∼ experience + abs. (lateralization index), family=Gamma, link=log. AIC = 52.4. Nagelkerke’s R2 = 0.127. Gamma dispersion = 0.407. Log estimates are listed below.

|  | *Estimate* | *Std. Error* | *Z value* | *Pr (>\|z\|)* |
| --- | --- | --- | --- | --- |
| **Intercept** | **-0.6403** | **0.1721** | **-3.722** | **0.000449\*\*\*** |
| **Experience: N** | **0.4690** | **0.1853** | **2.531** | **0.014120\*** |
| abs. (LI) | 0.2264 | 0.2945 | 0.769 | 0.445134 |

**Table S4B.** – **Backtransforned estinates and G5% CI for Model S4A** using the exponential function.

|  | *Estimate* | *2.5%* | *97.5%* |
| --- | --- | --- | --- |
| Intercept | 0.527115 | 0.3843262 | 0.7378704 |
| Experience: N | 1.5983532 | 1.1162862 | 2.3232028 |
| abs. (LI) | 1.2540736 | 0.7332799 | 2.1595933 |

**Tables S5.** - **Robust Linear Mixed Model** for the average bout position of three body parts for each of the experienced moths. Formula: bout average ∼ 0 + body part * bias direction + (1 | animal/bout). Observation count = 6,828. Random-effects groups: animal=45, bout:animal=2,276. Inclusion of 0 as fixed factor removes overall intercept; instead, model reports per-combination estimated mean.

| *Group* | *Effect* | *Variance* | *SD* |
| --- | --- | --- | --- |
| Bout:animal | Intercept | 29.690 | 5.449 |
| Animal | Intercept | 1.227 | 1.108 |
| Residual | - | 6.618 | 2.573 |

**Table S5A.** *–* Random effects for robust mixed effect model, including weights applied to residuals and random-effects for outlier downweighting. Bout:animal considers nested structure of multiple points measured per bout, per animal.

**Residual robustness weights**
| *Metric* | *Value* |
| --- | --- |
| ~1 weights | 5,784 |
| Remaining weights (min. / Q1 / median / mean / Q3 / max.) | 0.217 / 0.640 / 0.819 / 0.771 / 0.935 / 0.999 |

**Random-effects robustness weights**
| *Metric* | *Value* |
| --- | --- |
| ~1 weights | 1,844 |
| Remaining weights (min. / Q1 / median / mean / Q3 / max.) | 0.255 / 0.604 / 0.758 / 0.736 / 0.903 / 0.998 |

**Table S5B.**
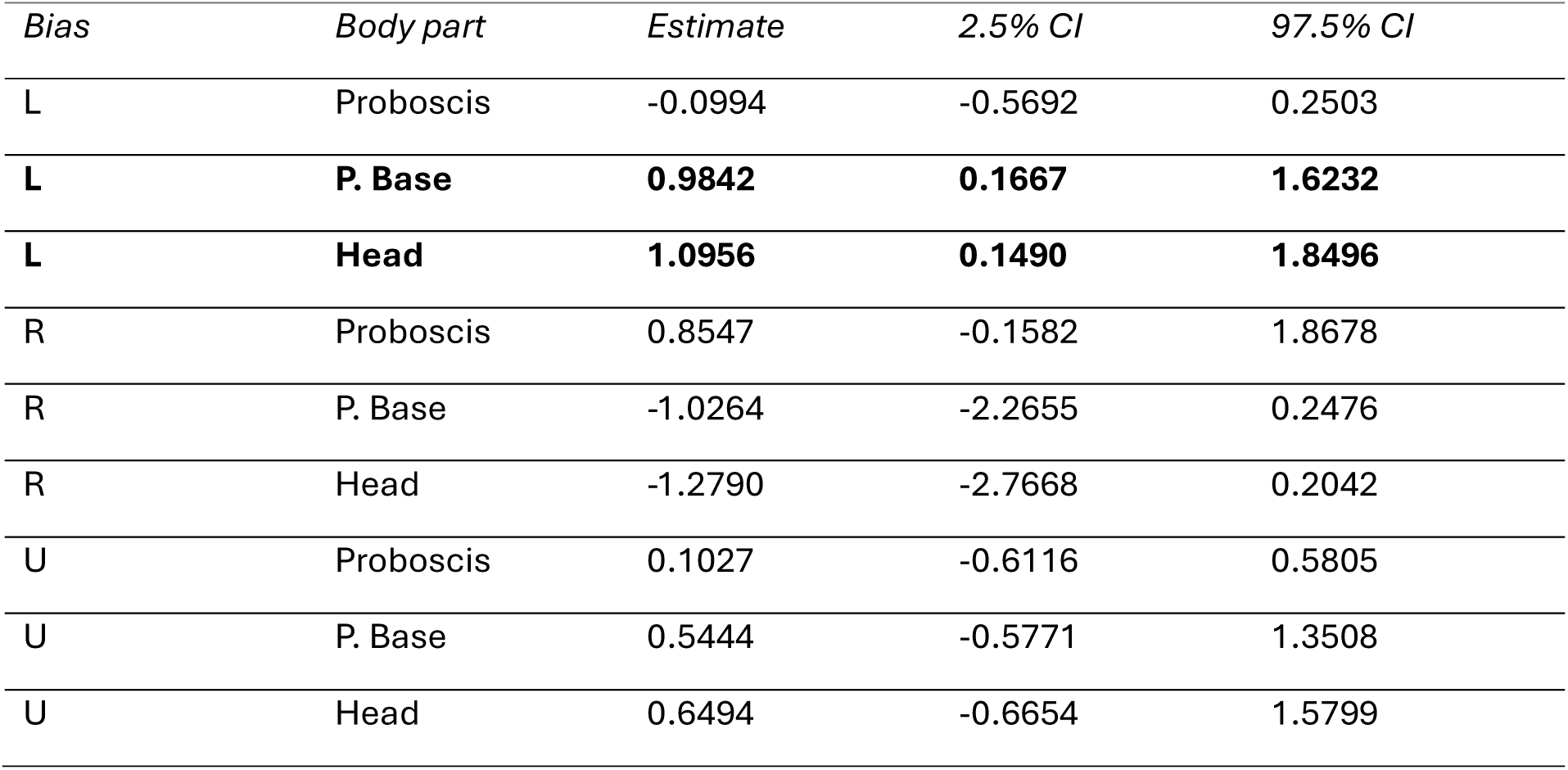
– Robust mixed model cell means by bias * body part. Estimates represent average position for respective body part (mm), relative to pattern midline. 95% confidence intervals for the estimate were obtained from 5000 cluster bootstrap resamples of model fit. Confidence intervals indicating the presence of a significant effect are in bold.

| *Bias* | *Body part* | *Estimate* | *2.5% CI* | *97.5% CI* |
| --- | --- | --- | --- | --- |
| L | Proboscis | -0.0994 | -0.5692 | 0.2503 |
| **L** | **P. Base** | **0.9842** | **0.1667** | **1.6232** |
| **L** | **Head** | **1.0956** | **0.1490** | **1.8496** |
| R | Proboscis | 0.8547 | -0.1582 | 1.8678 |
| R | P. Base | -1.0264 | -2.2655 | 0.2476 |
| R | Head | -1.2790 | -2.7668 | 0.2042 |
| U | Proboscis | 0.1027 | -0.6116 | 0.5805 |
| U | P. Base | 0.5444 | -0.5771 | 1.3508 |
| U | Head | 0.6494 | -0.6654 | 1.5799 |

**Table S5C.** – Robust mixed model within-bias body part contrasts. Estimates indicate difference between former and latter body part estimates. Confidence intervals indicating the presence of a significant effect are in bold.

| *Bias* | *Contrast* | *Estimate* | *2.5% CI* | *97.5% CI* |
| --- | --- | --- | --- | --- |
| **L** | **P. Base – Prob.** | **1.0835** | **0.6099** | **1.5485** |
| **L** | **Head – Prob.** | **1.1950** | **0.5976** | **1.7864** |
| L | Head – P. Base | 0.1115 | -0.0229 | 0.2401 |
| **R** | **P. Base – Prob.** | **-1.8811** | **-3.0632** | **-0.7042** |
| **R** | **Head – Prob.** | **-2.1337** | **-3.5056** | **-0.7406** |
| **R** | **Head – P. Base** | **-0.2526** | **-0.5057** | **-0.0124** |
| U | P. Base – Prob. | 0.4417 | -0.1328 | 0.9029 |
| U | Head – Prob. | 0.5468 | -0.2175 | 1.1347 |
| U | Head – P. Base | 0.1050 | -0.0899 | 0.2476 |

**Table S5D.**
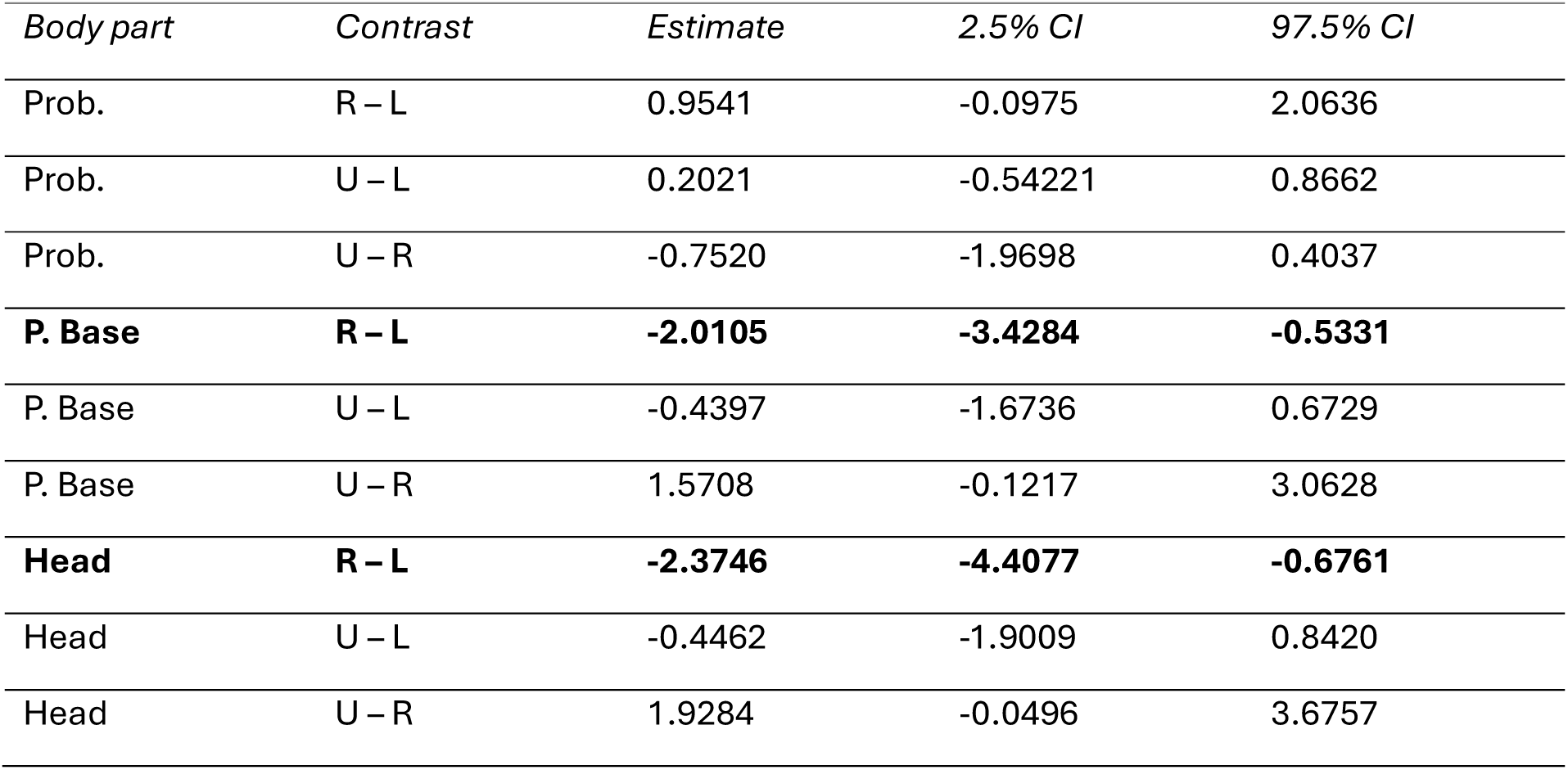
– Robust mixed model between-bias body part contrasts. Estimates indicate difference between estimated body part position between bias groups. Confidence intervals indicating the presence of a significant effect are in bold.

**Tables S6.** - **Robust Linear Mixed Model** for the average bout position of three body parts for each of the experienced moths used in monocular occlusion experiment. Formula: bout average ∼ 0 + body part * eye condition + (1 | animal/bout). Observation count = 4,476. Random-effects groups: animal=21, bout:animal=1,492. Inclusion of 0 as fixed factor removes overall intercept; instead model reports per-combination estimated mean. *Table S4A* *–* Random effects for robust mixed effect model, including weights applied to residuals and random-effects for outlier downweighting. Bout:animal considers nested structure of multiple points measured per bout, per animal.

| *Group* | *Effect* | *Variance* | *SD* |
| --- | --- | --- | --- |
| Bout:animal | Intercept | 27.614 | 5.255 |
| Animal | Intercept | 2.974 | 1.725 |
| Residual | - | 5.756 | 2.399 |

Residual robustness weights
| *Metric* | *Value* |
| --- | --- |
| ~1 weights | 3,826 |
| Remaining weights (min. / Q1 / median / mean / Q3 / max.) | 0.183 / 0.653 / 0.818 / 0.776 / 0.947 / 0.999 |

**Random-effects robustness weights**
| *Metric* | *Value* |
| --- | --- |
| ~1 weights | 1,225 |
| Remaining weights (min. / Q1 / median / mean / Q3 / max.) | 0.211 / 0.574 / 0.758 / 0.730 / 0.914 / 0.999 |

**Table S6B.** – Robust mixed model cell means by eye condition * body part. Estimates represent average position for respective body part (mm), relative to pattern midline. 95% confidence intervals for the estimate were obtained from 5000 cluster bootstrap resamples of model fit. LU = left bias, left eye unpainted; LP = left bias, left eye painted; RU = right bias, right eye unpainted; RP = right bias, right eye painted.

| *Eye Condition* | *Body part* | *Estimate* | *2.5% CI* | *97.5% CI* |
| --- | --- | --- | --- | --- |
| LU | Proboscis | -0.08135 | -0.93513 | 0.40835 |
| LU | P. Base | 0.96580 | -0.38690 | 1.77033 |
| LU | Head | 1.10276 | -0.43277 | 2.10080 |
| LP | Proboscis | -0.19778 | -1.00067 | 0.60191 |
| LP | P. Base | -0.15699 | -1.69157 | 1.48248 |
| LP | Head | -0.30178 | -2.19951 | 1.69587 |
| RU | Proboscis | 0.44281 | -0.29480 | 1.61501 |
| RU | P. Base | -1.63166 | -3.31514 | 0.62130 |
| RU | Head | -1.93343 | -3.97476 | 0.66243 |
| RP | Proboscis | 0.11471 | -1.69504 | 1.82659 |
| RP | P. Base | -0.11565 | -2.88161 | 2.25379 |
| RP | Head | 0.15840 | -2.83019 | 2.82121 |

**Table S6C.** – Robust mixed model within-condition body part contrasts. Estimates indicate difference between former and latter body part estimates. Confidence intervals indicating the presence of a significant effect are in bold.

| *Eye Condition* | *Contrast* | *Estimate* | *2.5% CI* | *97.5% CI* |
| --- | --- | --- | --- | --- |
| **LU** | **P. Base – Prob.** | **1.04714** | **0.41097** | **1.71228** |
| **LU** | **Head – Prob.** | **1.18411** | **0.34152** | **2.03088** |
| LU | Head – P. Base | 0.13697 | -0.07905 | 0.32808 |
| LP | P. Base – Prob. | 0.04078 | -1.15191 | 1.25173 |
| LP | Head – Prob. | -0.10399 | -1.64768 | 1.48852 |
| LP | Head – P. Base | -0.14478 | -0.52786 | 0.25442 |
| **RU** | **P. Base – Prob.** | **-2.07447** | **-3.51432** | **-0.65795** |
| **RU** | **Head – Prob.** | **-3.37624** | **-4.17192** | **-0.58768** |
| RU | Head – P. Base | -0.30177 | -0.67915 | 0.07883 |
| RP | P. Base – Prob. | -0.23036 | -1.25899 | 0.49564 |
| RP | Head – Prob. | 0.04369 | -1.26065 | 1.09276 |
| RP | Head – P. Base | 0.27406 | -0.07222 | 0.60872 |

**Table S6D.** – Robust mixed model of between-eye-condition body part contrasts. Estimates indicate difference between estimated body part position between bias groups.

| *Body part* | *Contrast* | *Estimate* | *2.5% CI* | *97.5% CI* |
| --- | --- | --- | --- | --- |
| Prob. | LP – LU | -0.11643 | -0.72379 | 0.83275 |
| Prob. | RP – RU | -0.32810 | -1.98334 | 0.50326 |
| P. Base | LP – LU | -1.12280 | -2.63330 | 0.63885 |
| P. Base | RP – RU | 1.151600 | -1.17075 | 3.22918 |
| Head | LP – LU | -1.40454 | -3.27456 | 0.72561 |
| Head | RP – RU | 2.09183 | -0.92789 | 4.01891 |

