## Supplementary Figures and Tables for "Conservation of a lateralized visuo-motor axis in hawkmoth proboscis probing"

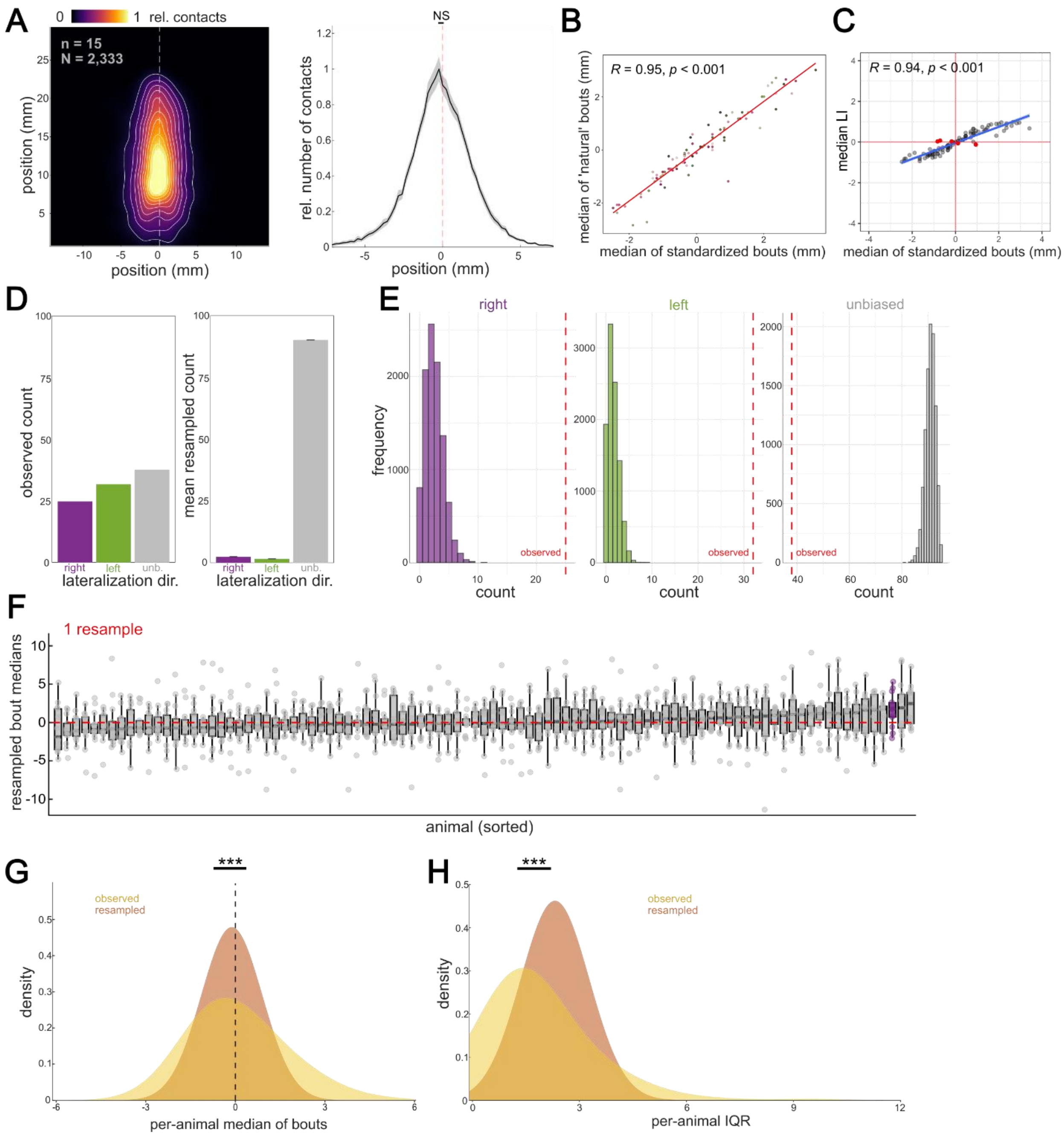

**Figure S1**

**A**: (left) Positions of the proboscis relative to the proboscis base during stimulus contact for unbiased hawkmoths, normalized and averaged across individuals (n) and bouts (N). (right) Normalized distribution of proboscis positions along the x-axis, indicating leftwards (negative) and rightwards (positive) proboscis positions. Distribution of per-animal medians was compared to zero using the Wilcoxon signed-rank test. **B:** Correlation between median proboscis lateralization amplitude of standardized bouts (see *Methods*) versus animal-generated bouts. Points depict individual hawkmoths. Red line depicts fitted linear regression. **C:** Correlation between median proboscis lateralization amplitude of standardized bouts and median lateralization index per animal. Points depict individual animals. Red points depict animals within incongruent quadrants (n=5). Blue line depicts fitted linear regression. **D:** Number of animals within each lateralization subgroup. Bars coloured by lateralization subgroup. (left) Counts observed in our data. (right) Mean count from resampled data (see *Methods*). Small black bars depict standard deviation of mean. **E:** Frequency of animals within each subgroup, per resampled iteration. Dashed red lines depict the number of animals observed in original data, per subgroup. **F:** Example distribution of lateralization amplitude bout maximums from one resampled iteration. Coloured distributions depict significant deviance from zero using Wilcoxon signed-rank test. **G-I:** (G) Density comparisons between observed (yellow) and resampled (brown) data for lateralization amplitude (G) and lateralization interquartile range (H). Comparisons between observed and resampled distributions were made using the two-sample Kolmogorov-Smirnov test. Statistical test results are abbreviated as * ($p<0.05$), ** ($p<0.01$), *** ($p<0.001$), and NS (not significant), with full details available in Tables S8-S12.

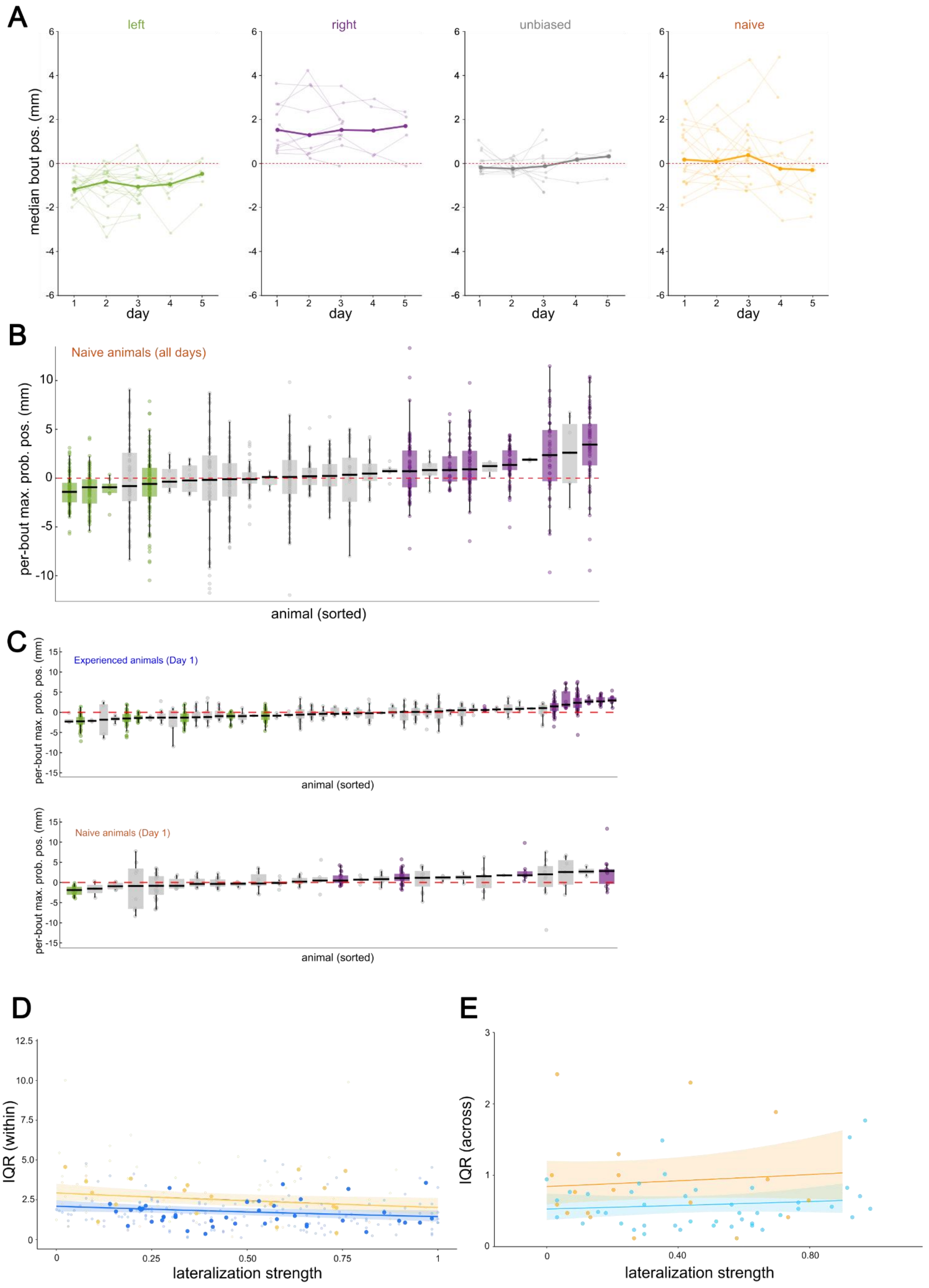

A
left
right
unbiased
naive
median bout pos. (mm)
day
B
Naive animals (all days)
per-bout max. prob. pos. (mm)
animal (sorted)
C
Experienced animals (Day 1)
per-bout max. prob. pos. (mm)
animal (sorted)
Naive animals (Day 1)
per-bout max. prob. pos. (mm)
animal (sorted)
D
IQR (within)
lateralization strength
E
IQR (across)
lateralization strength

**Figure S2**

**A**: Sequential medians of proboscis lateralization amplitude across days. Lines represent individual hawkmoths' median lateralization amplitude. Bolded lines depict median position of group per day. **B:** Distribution of bout-wise lateralization amplitudes for naïve hawkmoths across all days tested. Points and boxes are coloured by significance of median compared to zero using Wilcoxon signed-rank test. **C:** Distribution of bout-wise lateralization amplitudes for experienced (top) and naïve (bottom) hawkmoths on the first testing day. **D-E:** Fitted generalized linear mixed model for within-day interquartile range (D) as a function of lateralization amplitude (mm), and generalized linear model for across-day interquartile range (E), as a function of lateralization strength. (D) Large opaque points depict overall median within-day lateralization strength; small points depict median strength per day, coloured by experience group (naïve=yellow, blue=experienced). (E) Points depict overall median lateralization strength across days, per animal. Shaded bands depict 95% CI of fit. See Tables S3-S4 for statistical details of fitted models.

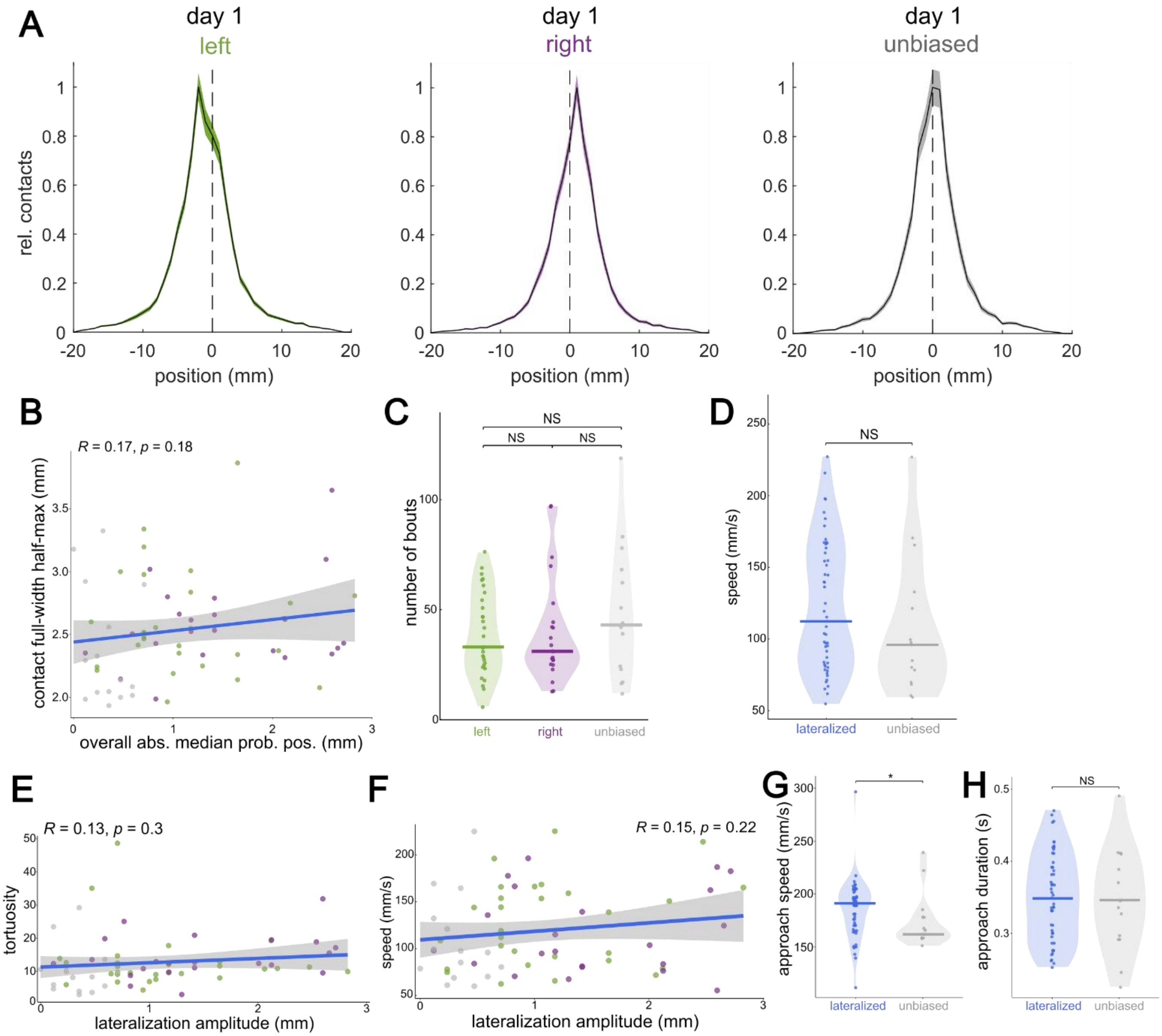

**Figure S3**

**A**: Normalized distributions of contact positions relative to the stripe midline (midline=0mm; edges=+/-1mm), on the first testing day, for each hawkmoth subgroup. **B:** Correlation between median absolute proboscis lateralization amplitude (mm) and full width at half maximum of the contact distribution (precision). Points indicate individual hawkmoths, coloured by lateralization subgroup. Blue line depicts fitted linear regression, shaded bands depict 95% CI. **C:** Number of bouts per lateralization subgroup. Points depict individual animals. Horizontal bars depict group median. Group distributions were compared with the Kruskal-Wallis test. **D:** Median bout-wise proboscis probing speed (mm/sec) between lateralized (blue) and unbiased (gray) hawkmoths. Group distributions were compared with the Wilcoxon rank-sum test. **E-F:** Correlations between absolute lateralization amplitude (mm) and (E) trajectory tortuosity, (F) probing speed (mm/sec). Blue lines depict fitted linear regressions, shaded bands depict 95% CI. **G:** Comparison of median bout-wise approach speeds (mm/sec) for lateralized versus unbiased hawkmoths. Group distributions were compared with the Wilcoxon rank-sum test. **H:** Comparison of median bout-wise approach duration (sec) for lateralized versus unbiased hawkmoths. Group distributions were compared with the Wilcoxon rank-sum test. Statistical test results are abbreviated as * ($p<0.05$), ** ($p<0.01$), *** ($p<0.001$), and NS (not significant), with full details available in Tables S8-S12.

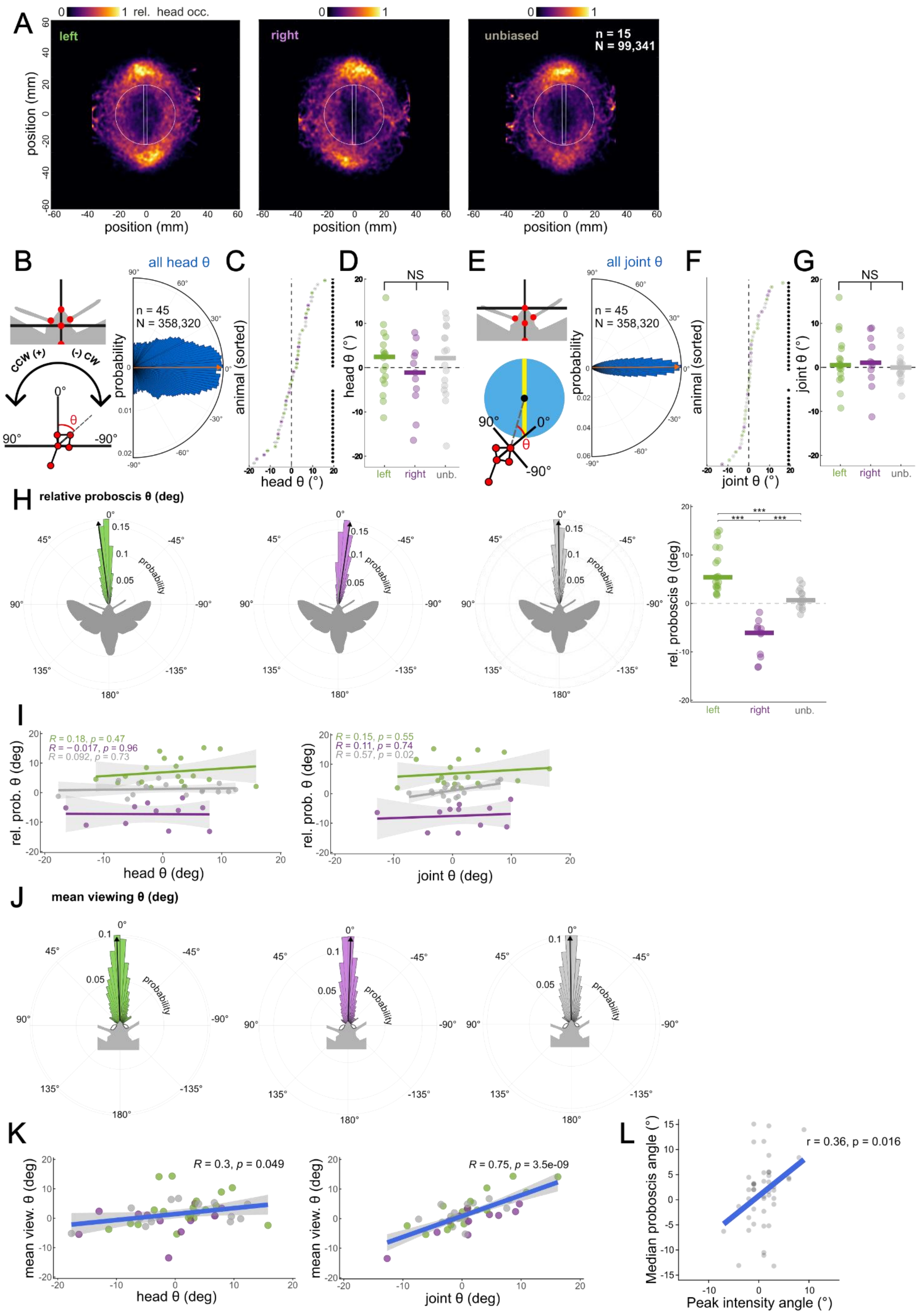
A
0 1 rel. head occ.
left
right
unbiased
n = 15
N = 99,341
position (mm)
position (mm)
B
all head θ
ccw (+)
(-) cw
n = 45
N = 358,320
probability
C
animal (sorted)
head θ (°)
D
NS
head θ (°)
left right unb.
E
all joint θ
n = 45
N = 358,320
probability
F
animal (sorted)
joint θ (°)
G
NS
joint θ (°)
left right unb.
H
relative proboscis θ (deg)
probability
rel. proboscis θ (deg)
***
left right unb.
I
R = 0.18, p = 0.47
R = −0.017, p = 0.96
R = 0.092, p = 0.73
rel. prob. θ (deg)
head θ (deg)
R = 0.15, p = 0.55
R = 0.11, p = 0.74
R = 0.57, p = 0.02
rel. prob. θ (deg)
joint θ (deg)
J
mean viewing θ (deg)
probability
K
R = 0.3, p = 0.049
mean view. θ (deg)
head θ (deg)
R = 0.75, p = 3.5e-09
mean view. θ (deg)
joint θ (deg)
L
r = 0.36, p = 0.016
Median proboscis angle (°)
Peak intensity angle (°)

**Figure S4:**

**A**: Normalized and averaged occupancy of the tracked head keypoints across all moths (n) and frames (N) where the proboscis was in contact, separated by subgroup (n/N for left and right subgroups in Fig. 4). White outline depicts the stimulus location. **B:** Distributions of head orientation during all proboscis contacts. Schematic (left) illustrates reference frame and polar coordinate system used. Coloured bars depict histogram probability. Orange arrow depicts overall median angle. **C:** Per-animal mean head orientation, sorted in ascending order. Points are coloured by lateralization group. Horizontal bars within points depict 95% CI of mean angle. Individual means tested for significance using the one-sample circular mean test. **D:** Mean head orientation angle for all lateralization subgroups. Points depict individual animals. Horizontal bars depict group medians. Comparisons were made using Kruskal-Wallis test. **E:** Distributions of joint angle between proboscis base and pattern centre during all proboscis contacts. Schematic (left) illustrates reference frame and polar coordinate system used. Coloured bars depict histogram probability. Orange arrow depicts overall median angle. **C:** Per-animal mean joint angle, sorted in ascending order. Points are coloured by lateralization group. Horizontal bars within points depict 95% CI of mean angle. Individual means tested for significance using the one-sample circular mean test. **D:** Mean joint orientation angle for all lateralization subgroups. Points depict individual animals. Horizontal bars depict group medians. Comparisons were made using Kruskal-Wallis test. **H:** (left) Circular distribution of relative proboscis angles during all contacts, sorted by lateralization subgroup. Coloured bars depict histogram probability. Black arrows depict median proboscis angle of group distributions. (right) Comparison of per-animal median proboscis angle between subgroups. Points depict individual animals, bars depict group median. Comparisons between groups were made using Kruskal-Wallis test, followed by pairwise Wilcoxon tests. **I:** Fitted linear regressions of the relationship between mean head angle (left) or mean joint angle (right) and median relative proboscis angle. Separate regressions were fitted per lateralization subgroup, depicted by colour. Points depict individual animals. **J:** (left) Circular distribution of mean viewing angle (see *Methods*) during all contacts, sorted by lateralization subgroup. Coloured bars depict histogram probability. Black arrows depict median viewing angle of group distributions. (right) Comparison of per-animal median viewing angle between subgroups. Points depict individual animals, bars depict group median. Comparisons between groups were made using Kruskal-Wallis test, followed by pairwise Wilcoxon tests. **K:** Fitted linear regressions of the relationship between mean head angle (left) or mean joint angle (right) and mean viewing angle. Separate regressions were fitted per lateralization subgroup, depicted by colour. Points depict individual animals. **L:** Relationship between visual angle of maximum visual scene intensity and median proboscis angle, per animal. Points depict individual animals. Blue line depicts fitted linear model. Statistical test results are abbreviated as * ($p<0.05$), ** ($p<0.01$), *** ($p<0.001$), and NS (not significant), with full details available in Tables S8-S12.

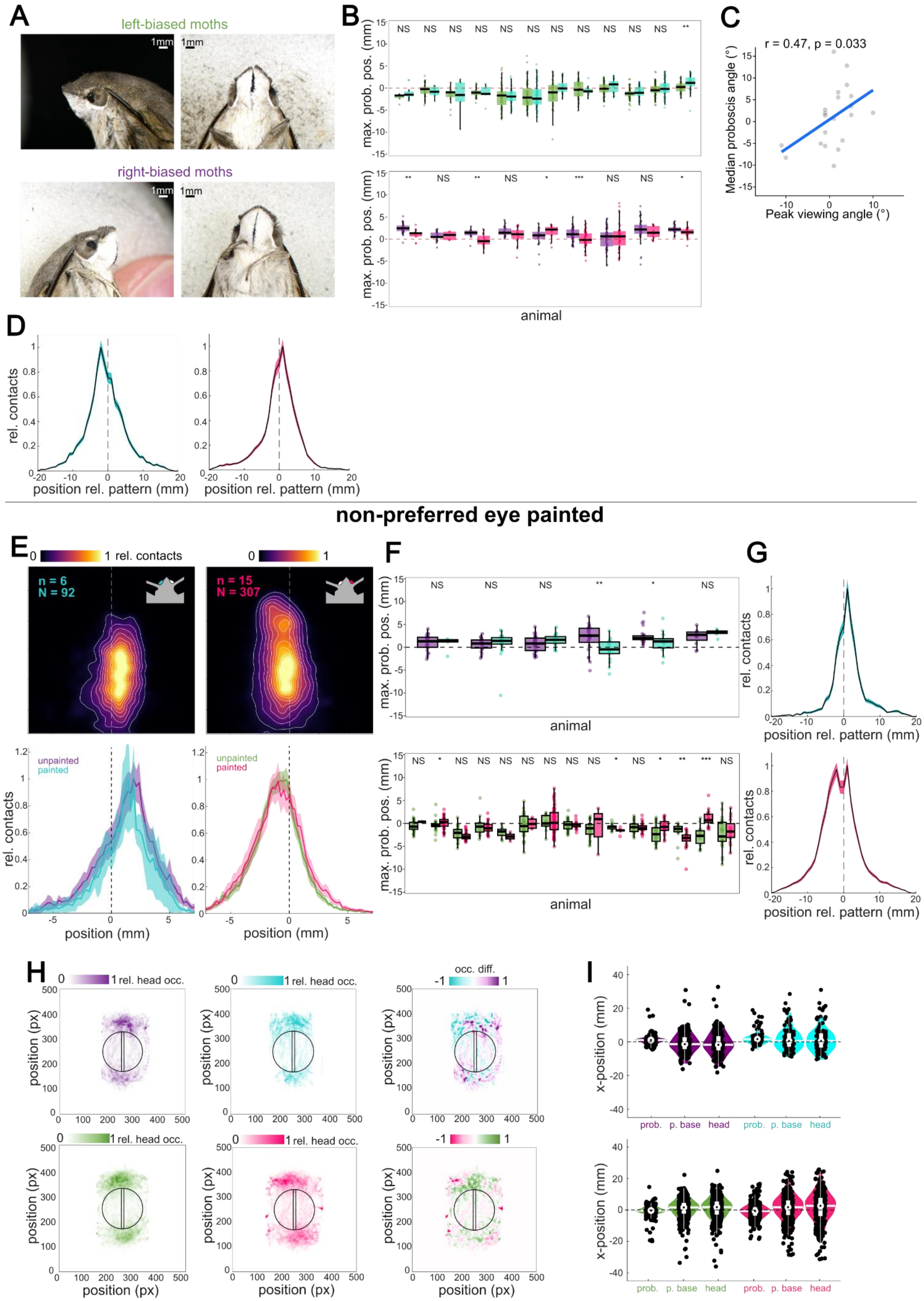
A
left-biased moths
right-biased moths
1mm
B
max. prob. pos. (mm)
NS
*
**
***
animal
C
r = 0.47, p = 0.033
Median proboscis angle (°)
Peak viewing angle (°)
D
rel. contacts
position rel. pattern (mm)
non-preferred eye painted
E
0
1 rel. contacts
n = 6
N = 92
n = 15
N = 307
unpainted
painted
rel. contacts
position (mm)
F
max. prob. pos. (mm)
animal
G
rel. contacts
position rel. pattern (mm)
H
1 rel. head occ.
occ. diff.
-1
position (px)
I
x-position (mm)
prob.
p. base
head

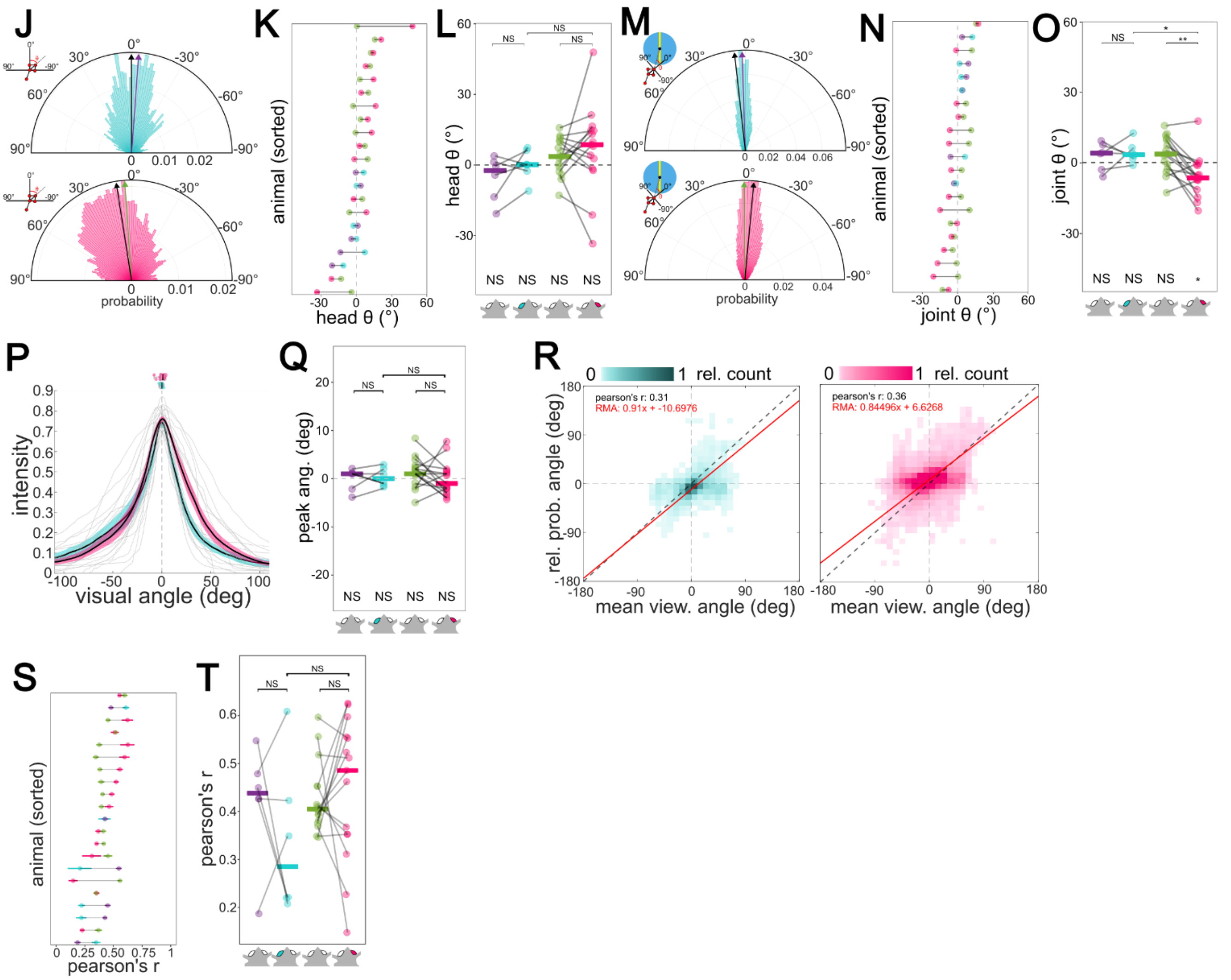

**Figure S5**

***Preferred eye painted:*** **A:** Images of left-lateralized (top) and right-lateralized (bottom) hawkmoths, with the fronto-ventral region of their respective eye painted. Inset bars depict 1mm scale. **B:** Distribution of per-animal lateralization amplitude maximums per bout, before (left of each column) and after (right of each column) eye painting (top=left eye, bottom=right eye). Bars depict medians, vertical lines depict whiskers. Each animal's distributions were compared before/after using paired Wilcoxon signed-rank tests. **C:** Relationship between visual angle of maximum visual scene intensity and median proboscis angle, per animal, following painting. Points depict individual animals. Blue line depicts fitted linear model. **D:** Normalized distributions of contact positions relative to the stripe midline (midline=0mm; edges=+/-1mm), for all data following painting.

***Non-preferred eye painted:*** **E:** (top) Positions of the proboscis relative to the proboscis base during stimulus contact for left- (left) and right-lateralized (right) hawkmoths following painting, normalized and averaged across individuals (n) and bouts (N). (bottom) Normalized distribution of proboscis positions along the x-axis. Before- and after-painting distributions are separated by colour. **F:** Distribution of per-animal lateralization amplitude maximums per bout, before (left of each column) and after (right of each column) eye painting (top=left eye, bottom=right eye). Bars depict medians, vertical lines depict whiskers. Each animal's distributions were compared before/after using paired Wilcoxon signed-rank tests. **G:** Normalized distributions of contact positions relative to the stripe midline (midline=0mm; edges=+/-1mm), for all data following painting. **H:** (left) Normalized and averaged difference heatmaps depicting the difference in head occupancy before and after painting for both lateralized subgroups (top: left eye, right-lateralized, bottom: right eye, left-lateralized). (right) Bout-wise average positions of tracked keypoints (mm) before and after painting, for both subgroups. **J:** Distributions of head orientation during all proboscis contacts following occlusion. Coloured bars depict histogram probability. Black arrows depict painted group median angle. Coloured arrows depict group median angle before painting. **K:** Per-animal mean head orientation, sorted in ascending order. Points are coloured by painted condition, with black lines depicting magnitude of change in angle. Horizontal bars within points depict 95% CI of mean angle. Individual means tested for significance using the one-sample circular mean test. **L:** Mean head orientation angle before and after painting, for both lateralization subgroups. Horizontal bars depict group medians. Within-group comparisons were made using Wilcoxon signed-rank, between-group comparisons were performed with Wilcoxon rank-sum. Group distributions were compared against zero using Wilcoxon signed-rank tests. **M:** Mean joint angle before and after painting. Within-group comparisons were made using Wilcoxon signed-rank, between-group comparisons were performed with Wilcoxon rank-sum. **N:** Per-animal mean joint angle, sorted in ascending order. Points are coloured by painted condition, with black lines depicting magnitude of change in angle. Horizontal bars within points depict 95% CI of mean angle. Individual means tested for significance using the one-sample circular mean test. **O:** Mean joint angles before and after painting, for both lateralization subgroups. Horizontal bars depict group medians. Within-group comparisons were made using Wilcoxon signed-rank, between-group comparisons were performed with Wilcoxon rank-sum. Group distributions were compared against zero using Wilcoxon signed-rank tests. **P:** Averaged visual scene intensity distributions. Boxplots depict distributions of per-animal angles of maximum intensity, separated by painted eye. **Q:** Comparison of maximum intensity angles before and after painting. Horizontal bars depict group medians. Within-group comparisons were made using Wilcoxon signed-rank, between-group comparisons were performed with Wilcoxon rank-sum. **R:** Density scatterplots of per-contact mean viewing angle and relative proboscis angles across all moths and contacts for the left-eye and right-eye painted groups, respectively. **S:** Pearson correlation strengths of per-contact mean viewing angle and relative proboscis angle, per animal. Animals sorted in order of ascending correlation strength. Colours of points depict painted group and eye condition. Black horizontal lines depict magnitude of change in coefficient. **T:** Pearson correlation strengths before and after monocular occlusion. Horizontal bars depict group median. Within-group comparisons were made using the Wilcoxon signed-rank test. Between-group comparisons were made using the Wilcoxon rank-sum test. Statistical test results are abbreviated as * ($p<0.05$), ** ($p<0.01$), *** ($p<0.001$), and NS (not significant), with full details available in Tables S8-S12.

Table S7 – Wilcoxon signed-rank tests (against zero; two-tailed) performed throughout the study

| *Figure* | *n* | *N* | *Test statistic* | *p-value* | *Corrected p (when applicable)* |
|---|---|---|---|---|---|
| 1 *F* | 95 | 95 | 2294.5 | 0.8166 | 0.8166 |
| **1 *H* (left)** | **32** | **32** | **0** | **8.247e-07** | **2.4741e-06** |
| **1 *H* (right)** | **25** | **25** | **325** | **1.304e-05** | **2.6080e-05** |
| S1 *A* | 38 | 38 | 409.5 | 0.3851 | 0.3851 |
| S4 *D* | L: 18<br>R: 11<br>U: 16 | L: 18<br>R: 11<br>U: 16 | L: 108<br>R: 27<br>U: 83 | L: 0.3465<br>R: 0.6377<br>U: 0.4637 | L: 0.3465<br>R: 0.6377<br>U: 0.4637 |
| S4 *G* | L: 18<br>R: 11<br>U: 16 | L: 18<br>R: 11<br>U: 16 | L: 103<br>R: 43<br>U: 67 | L: 0.4683<br>R: 0.4131<br>U: 0.9799 | L: 0.4683<br>R: 0.4131<br>U: 0.9799 |
| **S4 *H*** | **L: 18**<br>**R: 11**<br>U: 16 | **L: 18**<br>**R: 11**<br>U: 16 | **L: 190**<br>**R: 25**<br>U: 78.5 | **L: 0.000143**<br>**R: 0.003857**<br>U: 0.09384 | **L: 0.000429**<br>**R: 0.007714**<br>U: 0.09384 |
| 5 *F* | LU: 12<br>LP: 12<br>RU: 9<br>RP: 9 | LU: 12<br>LP: 12<br>RU: 9<br>RP: 9 | LU: 52<br>LP: 27<br>RU: 15<br>RP: 32 | LU: 0.3268<br>LP: 0.367<br>RU: 0.4069<br>RP: 0.2863 | LU: 0.3268<br>LP: 0.367<br>RU: 0.4069<br>RP: 0.2863 |
| **5 *I*** | LU: 12<br>**LP: 12**<br>RU: 9<br>**RP: 9** | LU: 12<br>**LP: 12**<br>RU: 9<br>**RP: 9** | LU: 39<br>**LP: 72**<br>RU: 32<br>**RP: 2** | LU: 1<br>**LP: 0.01079**<br>RU: 0.2863<br>**RP: 0.01782** | LU: 1<br>**LP: 0.03564**<br>RU: 0.57260<br>RP: 0.05346 |
| **5 *K*** | LU: 12<br>**LP: 12**<br>RU: 9<br>RP: 9 | LU: 12<br>**LP: 12**<br>RU: 9<br>RP: 9 | LU: 46.5<br>**LP: 70.5**<br>RU: 26<br>RP: 17.5 | LU: 0.5775<br>**LP: 0.01452**<br>RU: 0.7194<br>RP: 0.5903 | LU: 1<br>LP: 0.05808<br>RU: 0.71940<br>RP: 0.71940 |

| | | | | | |
|---|---|---|---|---|---|
| S5 *L* | LU: 6<br>LP: 6<br>RU: 15<br>RP: 15 | LU: 6<br>LP: 6<br>RU: 15<br>RP: 15 | LU: 5<br>LP: 12<br>RU: 87<br>RP: 91 | LU: 0.2945<br>LP: 0.8339<br>RU: 0.1323<br>RP: 0.08322 | LU: 0.2945<br>LP: 0.8339<br>RU: 0.1323<br>RP: 0.08322 |
| **S5 *O*** | LU: 6<br>LP: 6<br>RU: 15<br>**RP: 15** | LU: 6<br>LP: 6<br>RU: 15<br>**RP: 15** | LU: 16<br>LP: 20<br>RU: 87<br>**RP: 16** | LU: 0.2945<br>LP: 0.05917<br>RU: 0.1323<br>**RP: 0.01349** | LU: 0.29450<br>LP: 0.17751<br>RU: 0.25460<br>RP: 0.05396 |
| S5 *Q* | LU: 6<br>LP: 6<br>RU: 15<br>RP: 15 | LU: 6<br>LP: 6<br>RU: 15<br>RP: 15 | LU: 10.5<br>LP: 12.5<br>RU: 87.5<br>RP: 50.5 | LU: 1<br>LP: 0.7498<br>RU: 0.122<br>RP: 0.6069 | LU: 1<br>LP: 0.7498<br>RU: 0.122<br>RP: 0.6069 |

Table S8 – Wilcoxon signed-rank tests (paired; two-tailed) performed throughout the study

| *Figure* | *n* | *N* | *Test statistic* | *p-value* | *Corrected p* |
|---|---|---|---|---|---|
| 5 *F* | LU – LP: 12<br>RU – RP: 9 | LU – LP: 12<br>RU – RP: 9 | LU – LP: 57<br>RU – RP: 8 | LU – LP: 0.1698<br>RU – RP: 0.0972 | LU – LP: 0.1698<br>RU – RP: 0.0972 |
| **5 *I*** | **LU – LP: 12**<br>**RU – RP: 9** | **LU – LP: 12**<br>**RU – RP: 9** | **LU – LP: 10**<br>**RU – RP: 45** | **LU – LP: 0.02537**<br>**RU – RP: 0.009152** | **LU – LP: 0.02537**<br>**RU – RP: 0.018304** |
| *5 K* | LU – LP: 12<br>RU – RP: 9 | LU – LP: 12<br>RU – RP: 12 | LU – LP: 9<br>RU – RP: 27.5 | LU – LP: 0.2322<br>RU – RP: 0.2048 | LU – LP: 0.2322<br>RU – RP: 0.2048 |
| **5 *N*** | **LU – LP: 12**<br>**RU – RP: 9** | **LU – LP: 12**<br>**RU – RP: 9** | **LU – LP: 3**<br>**RU – RP: 0** | **LU – LP: 0.005355**<br>**RU – RP: 0.009152** | **LU – LP: 0.009152**<br>**RU – RP: 0.009152** |
| S5 *L* | LU – LP: 6<br>RU – RP: 15 | LU – LP: 6<br>RU – RP: 15 | LU – LP: 3<br>RU – RP: 49 | LU – LP: 0.1422<br>RU – RP: 0.5509 | LU – LP: 0.1422<br>RU – RP: 0.5509 |
| **S5 *O*** | LU – LP: 6<br>**RU – RP: 15** | LU – LP: 6<br>**RU – RP: 15** | LU – LP: 8<br>**RU – RP: 110** | LU – LP: 0.675<br>**RU – RP: 0.004932** | LU – LP: 0.675<br>**RU – RP: 0.009864** |

| | | | | | |
|---|---|---|---|---|---|
| S5 *Q* | LU – LP: 6<br>RU – RP: 15 | LU – LP: 6<br>RU – RP: 15 | LU – LP: 6<br>RU – RP: 75 | LU – LP: 0.7835<br>RU – RP: 0.1659 | LU – LP: 0.7835<br>RU – RP: 0.1659 |
| S5 *T* | LU – LP: 6<br>RU – RP: 15 | LU – LP: 6<br>RU – RP: 15 | LU – LP: 17<br>RU – RP: 40 | LU – LP: 0.2084<br>RU – RP: 0.2681 | LU – LP: 0.2084<br>RU – RP: 0.2681 |

Table S9 – Wilcoxon rank-sum tests (two-tailed) performed throughout the study

| *Figure* | *comparison* | *Test statistic* | *p-value* | *Corrected p-value* |
|---|---|---|---|---|
| **1 *H*** | **Left vs. right contact dists.** | **0** | **1.3e-10** | - |
| S1 *F* | Original vs. resampled medians | 4451.5 | 0.8642 | - |
| **S1 *G*** | **Original vs. resampled IQR** | **2557.5** | **1.943e-07** | - |
| **3 E** | **Lateralized vs. unbiased FWHM** | **521** | **0.03472** | - |
| 3 H | Lateralized vs. unbiased duration | 507 | 0.05778 | - |
| **3 G** | **Lateralized vs. unbiased length** | **574** | **0.003472** | **-** |
| **3 F** | **Lateralized vs. unbiased tortuosity** | **527** | **0.02757** | **-** |
| S3 D | Lateralized vs. unbiased probing speed | 468 | 0.1934 | - |
| **S3 G** | **Lateralized vs. unbiased approach speed** | **530** | **0.02447** | **-** |
| S3 H | Lateralized vs. unbiased approach duration | 401.5 | 0.777 | - |

| | | | | |
|---|---|---|---|---|
| **4 *F*** | **Median viewing angle (left vs. right experienced)** | **155.5** | **0.02863** | **-** |
| 5 *F* | Mean head angle (left painted vs. right painted) | 31 | 0.1098 | - |
| **5 *I*** | **Mean joint angle (left painted vs. right painted)** | **99** | **0.001564** | **4.692e-03** |
| **5 *K*** | **Peak viewing angle (left painted vs. right painted)** | **83** | **0.04061** | 0.12183 |
| S5 *L* | Mean head angle (left contralateral painted vs. right contralateral painted) | 24 | 0.1105 | - |
| **S5 *O*** | **Mean joint angle (left contralateral painted vs. right contralateral painted)** | **81** | **0.005716** | **0.017148** |
| S5 *Q* | Peak viewing angle (left contralateral painted vs. right contralateral painted) | 55 | 0.4551 | 0.4551 |

Table S10 – Two-sample Kolmogorov-Smirnov tests performed throughout the study

| *Figure* | *comparison* | *Test statistic* | *p-value* |
|---|---|---|---|
| **S1 *F*** | **Original vs. resampled median dists.** | **0.48421** | **2.33e-16** |

| | | | |
|---|---|---|---|
| **S1 *G*** | **Original vs. resampled IQR dists.** | **0.70526** | **< 2.2e-16** |

Table S11 – Kruskal-Wallis tests performed throughout the study

| *Figure* | *Comparison* | *df* | *Test statistic* | *p-value* | *Pairwise wilcox test with Hommel correction* |
|---|---|---|---|---|---|
| **3 *B*** | **Maximum proboscis position (L/R/U)** | **2** | **7.8416** | **0.01982** | **r-l : 0.029**<br>r-u: 0.398<br>l-u: 0.109 |
| 3 *D* | Abs. prob.-edge distance (mm) (L/R/U) | 2 | 1.865 | 0.3936 | 0.3936 |
| S3 *C* | Median number of bouts (L/R/U) | 2 | 0.63747 | 0.7271 | 0.7271 |
| S4 D | Mean head angle | 2 | 1.2492 | 0.5355 | 0.5355 |
| S4 *G* | mean joint head angle (L/R/U) | 2 | 0.83792 | 0.6577 | 0.6577 |
| 4 *I* | Pearson's r (L/R/U) | 2 | 2.7608 | 0.2515 | 0.2515 |
| **S4 *H*** | **Median rel. proboscis angles (L/R/U)** | **2** | **33.447** | **5.46e-08** | **r-l: 1.1e-07**<br>**r-u: 1.0e-06**<br>**l-u: 1.5e-05** |

Table S12 – Circular statistics using the circ_stat Matlab toolbox

| *Figure* | *Group* | *n* | *N* | *Test* | *Output* | *Stat. comp.* | *p-value* |
|---|---|---|---|---|---|---|---|
| S4 *B* | All experienced | 45 | 358,320 | One-sample | 0.25° | One-sample | 0.9614 |

| | | | | | | | |
|---|---|---|---|---|---|---|---|
| | moths head angles | | | circular median | | median test | |
| S4 *E* | All experienced moths joint angles | 45 | 358,320 | One-sample circular median | 0.596° | One-sample median test | 1 |
| 5 *D* | Ipsilateral LP head angles | 45 | 43,865 | One-sample circular median | -3.81° | One-sample median test | 1 |
| 5 *D* | Ipsilateral RP head angles | 45 | 34,040 | One-sample circular median | 8.83° | One-sample median test | 1 |
| 5 *G* | Ipsilateral LP joint angles | 45 | 43,865 | One-sample circular median | 4.16° | One-sample median test | 1 |
| 5 *G* | Ipsilateral RP joint angles | 45 | 34,040 | One-sample circular median | -4.39° | One-sample median test | 1 |
| **5 *K*** | **Ipsilateral LP viewing angles** | **12** | **12** | **Rayleigh test** | **mean = 3.59°, R = 0.994, circular SD = 6.29°** | **Rayleigh test of non-uniformity** | **2.452e-08** |
| **5 *K*** | **Ipsilateral LP viewing angles** | **12** | **12** | **V-test** | **-** | **V test of non-uniformity clustered about 0°** | **6.234e-07** |
| **5 *K*** | **Ipsilateral RP viewing angles** | **9** | **9** | **Rayleigh test** | **mean = -3.11°, R = 0.981, circular SD = 11.36°** | **Rayleigh test of non-uniformity** | **6.367e-06** |

| | | | | | | | |
|---|---|---|---|---|---|---|---|
| **5 *K*** | **Ipsilateral RP viewing angles** | **9** | **9** | **V-test** | **-** | **V test of non-uniformity clustered about 0°** | **1.634e-05** |
| S5 *J* | Contralateral LP head angles | 6 | 10,554 | One-sample circular median | -0.99° | One-sample median test | 1 |
| S5 *J* | Contralateral RP head angles | 15 | 53,267 | One-sample circular median | 12.81° | One-sample median test | 1 |
| S5 *M* | Contralateral LP joint angles | 6 | 10,554 | One-sample circular median | 3.36° | One-sample median test | 1 |
| S5 *M* | Contralateral RP joint angles | 15 | 53,267 | One-sample circular median | -5.18° | One-sample median test | 1 |
| **S5 *P*** | **Contralateral LP viewing angles** | **6** | **6** | **Rayleigh test** | **mean = 6.27°, R = 0.992, circular SD = 7.37°** | **Rayleigh test of non-uniformity** | **0.0004** |
| **S5 *P*** | **Contralateral LP viewing angles** | **6** | **6** | **V-test** | **-** | **V test of non-uniformity clustered about 0°** | **0.0003** |
| **S5 *P*** | **Contralateral RP viewing angles** | **15** | **15** | **Rayleigh test** | **mean = -1.37°, R = 0.982, circular SD = 10.91°** | **Rayleigh test of non-uniformity** | **5.316e-10** |
| **S5 *P*** | **Contralateral RP viewing angles** | **15** | **15** | **V-test** | **-** | **V test of non-uniformity clustered about 0°** | **3.779e-08** |
